# Antarctic biodiversity predictions through substrate qualities and environmental DNA

**DOI:** 10.1101/2021.08.18.456862

**Authors:** Paul Czechowski, Michel de Lange, Micheal Knapp, Aleks Terauds, Mark I. Stevens

## Abstract

Antarctic conservation science is important to enhance Antarctic policy and to understand alterations of terrestrial Antarctic biodiversity. Antarctic conservation will have limited long term effect in the absence of large-scale biodiversity data, but if such data were available, it is likely to improve environmental protection regimes. To enable Antarctic biodiversity prediction across continental spatial scales through proxy variables, in the absence of baseline surveys, we link Antarctic substrate-derived environmental DNA (eDNA) sequence data from the remote Antarctic Prince Charles Mountains to a selected range of concomitantly collected measurements of substrate properties. We achieve this using a statistical method commonly used in machine learning. We find neutral substrate pH, low conductivity, and some substrate minerals to be important predictors of presence for basidiomycetes, chlorophytes, ciliophorans, nematodes, or tardigrades. Our bootstrapped regression reveals how variations of the identified substrate parameters influence probabilities of detecting eukaryote phyla across vast and remote areas of Antarctica. We believe that our work may improve future taxon distribution modelling and aid targeting logistically challenging biodiversity surveys.

## Introduction

Although only 0.3% of continental Antarctica is ice-free, many organisms including bacteria, unicellular eukaryotes, fungi, lichen, cryptogamic plants and invertebrates are scattered across the continent in extremely isolated, remote, island-like terrestrial habitats, for example in soil-like substrates, lakes, and cryoconite holes (Convey *et al.* 2014; Chown *et al.* 2015). Threats to this Antarctic biodiversity are posed by human activity, climate change, pollution, and invasive species. It is becoming increasingly clear that mitigation of these threats and further alterations to the Antarctic biosphere rely on well-tailored management strategies across the continent’s bioregions (eg Coetzee *et al.* 2017).

Effective continental-scale conservation management requires continental-scale data (eg Wauchope *et al.* 2019). However, knowledge of terrestrial Antarctic biodiversity is still limited because most of Antarctica’s ice-free areas remain unstudied due to logistic difficulties exacerbated by the harsh environmental conditions, and funding constraints. Environmental DNA (eDNA) analysis, despite shortcomings, is arguably one of the most practical and economical options for continental-wide baseline surveys of terrestrial Antarctic biodiversity, especially when facing logistical challenges typical for work on the Antarctic continent (reviewed in Czechowski *et al.* 2017). Comparable large-scale systematic approaches to protect soil diversity are recognized as required globally, but often are limited to charismatic groups such as those found in the Arctic (Gillespie *et al.* 2020).

Here, we link commonly measured substrate properties to the cryptic eukaryotic biodiversity of terrestrial Antarctic ice-free regions. Soil nutrient status is the most important attribute of biodiverse soils (Geisen *et al.* 2019), and corresponding key variables can be, and are, routinely measured economically. We analyzed molecular data (eDNA) from an extremely remote Antarctic terrestrial region to clarify relationships between substrate properties and eukaryote phylum presence. We envisage our approach to be useful in predicting biodiversity across a wide taxonomic spectrum across large areas of Antarctica, especially to identify regions worthy of lower-level taxonomic biodiversity surveys, then possibly realized with “barcoding” using mitochondrial DNA (such as with the Cytochrome Oxidase 1) or logistically more challenging morphological biodiversity assessments.

The Prince Charles Mountains (PCMs), the most remote terrestrial areas in eastern Antarctica, were first sighted by US Operation Highjump (1946/47) and mapped in more detail by Australian (1954–1961) and Russian (1983–1991) expeditioners. In 2011 we obtained environmental DNA samples from substrates throughout the PCMs and measured various geochemical and mineral properties. Previously, Czechowski *et al.* (2016b) focused on invertebrates as the primary substrate-inhabiting metazoans and discovered major changes in their distribution over salinity gradients, as known from other areas and taxa of Antarctica (eg Bottos *et al.* 2020). Here, we expand our analyses of environmental variables using a predictive approach to the full spectrum of eukaryote phyla, and thereby explore approaches of inferring biodiversity presence that could be applied across the entirety of ice-free terrestrial Antarctica. Beyond phylum-level surveys, our technique may be applied using other genetic markers and predictors to link future smaller-scaled conservation projects anywhere in terrestrial Antarctica, aid taxon distribution modelling, and thus contributes towards improving conservation management strategies across the Antarctic bioregions.

## Methods

Fieldwork took place in the Prince Charles Mountains (PCMs; East Antarctica, Figure 1) from 26 November 2011 to 21 January 2012 close to Mount Menzies (MM; 73°25′29.38“S, 62°0′37.61“E), Mawson Escarpment (ME; 73°19′16.91“S, 68°19′31.20“E) and Lake Terrasovoje (LT; 70°32′23.58“S, 67°57′28.05“E) as described earlier (Czechowski *et al.* 2016a, b). 154 field samples were considered for this study (26 MM, 70 ME, 58 LT; Web Table 1).

**Figure 1:**
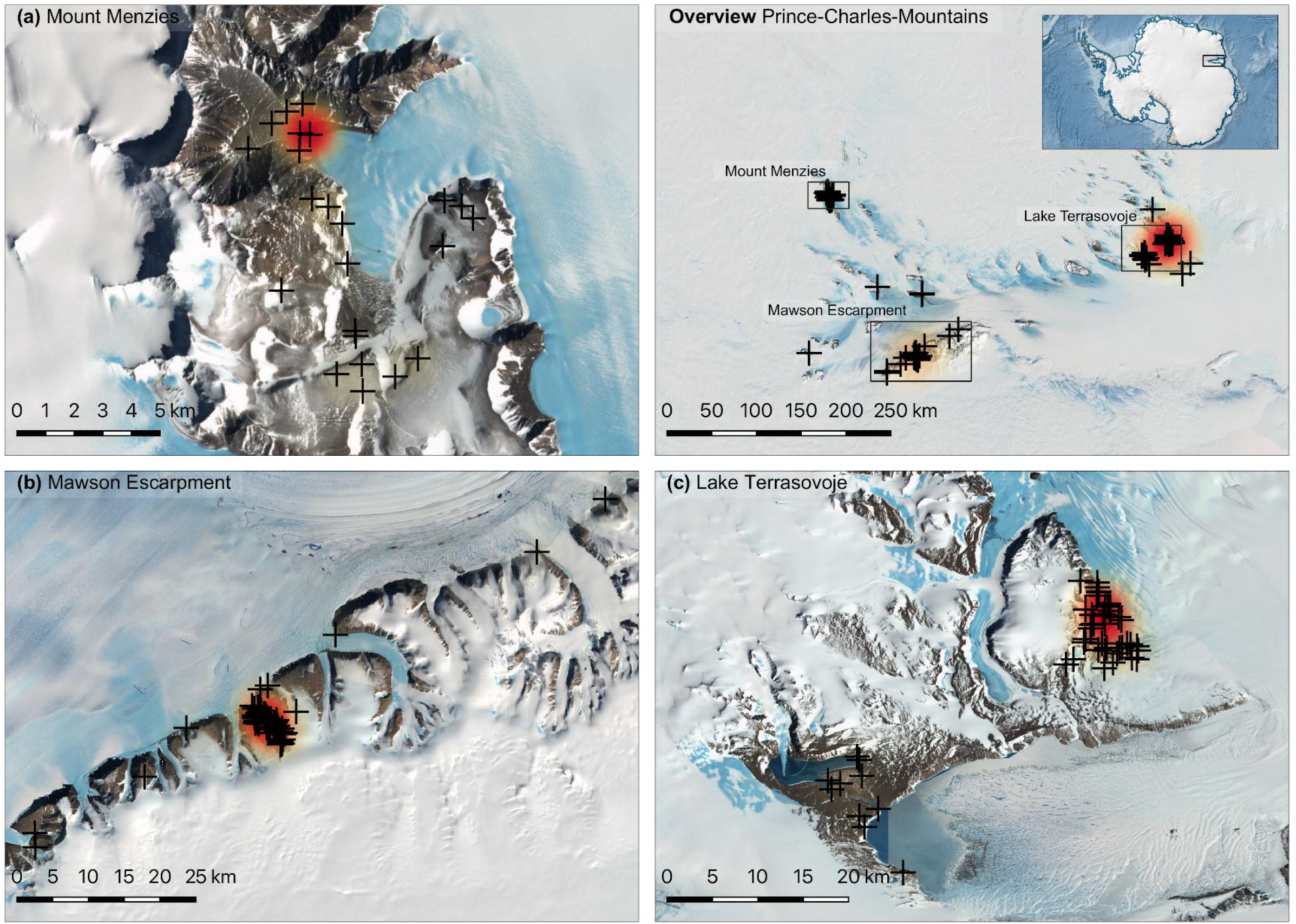
Sampling area. All sampling locations are marked with a crosshair. Heat shading (at map scale) indicates density of 18S Amplicon Sequence Variants (*sensu* Callahan *et al.* 2017) determined to be significantly influenced by substrate qualities as available. Base layers compiled by the Norwegian Polar Institute and distributed in the Quantarctica package. Visit http://www.quantarctica.org/. Base layers courtesy of the SCAR Antarctic Digital Database, © 1993–2015 Scientific Committee on Antarctic Research; The National Snow and Ice Data Centre, University of Colorado, Boulder; NASA, Visible Earth Team, http://visibleearth.nasa.gov/; Australian Antarctic Division, © Commonwealth of Australia 2006.

To infer climatic conditions in the PCMs, we used rasters from Quantarctica 3 (Matsuoka *et al.* 2021) encoding annual mean precipitation (mm), wind speed (m s^−1^ 10m above ground) and mean annual temperature (°C 2 m above ground, as only temperature data distributed via Quantarctica). We disaggregated the layer rasterization from 35 km px^−1^ to 1 km px^−1^ through bilinear interpolation. We then extracted median values for the three variables from a 20 km buffer surrounding each sampling location (Web Figure 1).

As predictor data for eukaryote phylum presence in substrates, geochemical composition (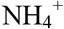, C, ρ, 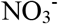, pH_H20_, pH_CaCl2_, P, K, S, texture) was analyzed by agricultural soil testing service APAL (www.apal.com.au). Many measurements below detection level needed to be excluded to yield data completeness of at least 96.7% (Web Table 2). The final analysis included K, S, ρ, and pH _CaCl2_ (pH_H2O_ excluded as co-linear, texture excluded as categorical). As additional predictors, the substrate mineral composition was considered through integration of X-ray diffraction spectra of the minerals quartz, calcite, feldspar, titanite, pyroxene / amphibole / garnet, micas, dolomite and kaolin / chlorite, and chlorite (see Czechowski *et al.* 2016b).We handled the sum-to-unity constraint of our mineral compositions by excluding quartz as the most common mineral from further analysis. As further predictors for most locations (MM: n=26, ME: n=69, LT: n=57), we included hitherto unpublished measurements of soil-substrate ATP (eg Conklin and Macgregor 1972), obtained with a Clean-Trace Luminometer (3M, Maplewood, US-MN), and slope measurements. Prior to regression, all predictors were standardized to mean of 0 and unit variance. Predictor densities are provided in Web Figure 2.

Biological response data were prepared in QIIME 2020-2 (Bolyen *et al.* 2019) and R 4.0.0 (R Core Development Team 2019) from raw sequence data generated as described elsewhere (Czechowski *et al.* 2016b, 2017). In summary, 125 bp eukaryotic 18S rDNA PCR products (yielding an 85 bp target region) had been amplified using primers ‘Euk1391f’ and ‘EukBr’ (Caporaso *et al.* 2012), as established for eukaryotic microbial surveying (Thompson *et al.* 2017). As recommended, PCRs had been carried out in triplicates, each replicate carrying identical barcodes. The resulting eDNA libraries had been combined for sequencing across two MiSeq runs (Web Figure 3). We re-defined Amplicon Sequence Variants (ASVs; *sensu* Callahan *et al.* 2017) from those data with Qiime: after pre-filtering (Phred score ≥ 25), we trimmed read pairs with Cutadapt v1.18 (Martin 2011), and denoised using DADA2 (v1.6.0; Callahan *et al.* 2016). We retained merged reads with an expected error value less than 3, that we not deemed chimeric.

Due to the shortness and slow evolution of the employed 18S marker, we set out to conduct our analyses on the phylum level, and to use species level assignments solely to verify data credibility. Accordingly, we designed the retrieval of taxonomic annotations for our Antarctic DNA sequences in such fashion so as to yield reliable species identifications in cases where Antarctic reference data were available, while still retuning higher taxonomic (eg phylum level) identifications in cases where closely matching reference data were not available. Doing so, we were able include a larger amount of Antarctic sequences into our statistical analysis on phylum level, but needed to consider species level identifications as potentially unreliable, and verify them on alignment level. We identified eukaryotic sequences among our reads with a recent local copy (April 2020) of the entire NCBI nucleotide collection in conjunction with Blast 2.10.0+. Taxonomic assignments were retrieved from reference sequences *at least* 50% identical to queries, with an assignment significance threshold (*e* value) of 10^−10^, considering only matches with at least 90% coverage, and excluding environmental sequences *(evalue 1e^−10^, max_hsps 5, max_target_seqs 5, qcov_hsp_perc 90* and *perc_identity 50)*. For each Antarctic sequence search query, we used the highest Bit score among all returned sequences from the NCBI database for that query to choose the final taxonomic assignment. Subsequently, we used R package *decontam* (Davis *et al.* 2018) to remove putatively contaminating reads, and likewise subtracted all sequences and taxa in negative controls from field samples. Focusing on eukaryotes, we discarded all non eukaryote reads (Web Figure 4).

With the Lasso (Tibshirani 1996) of R package *glmnet* (Friedman *et al.* 2010) we regressed each phylum present in at least 12 samples against the aforementioned predictors (Web Figure 5). In regressions, we disregarded sequence read abundances as meaningless due to inherent constraints of amplicon sequencing (eg Czechowski *et al.* 2017), analyzed presences instead, and used the most biodiverse of all locations (LT; Czechowski *et al.* 2016b; also Figure 2) as a reference location, so that we report predictor effects at MM and ME as relative to LT. We initially retrieved the active set (variables not set to 0) estimated by Lasso, repeated the regression of phylum presence against 1,000 randomly chosen sample-sets of predictors, calculated the number of times each variable was estimated to be non-zero, and report variables non-zero more than 950 times as significant. Accordingly, we calculated 95% non-parametric bootstrap confidence intervals for our estimates. We did not adjust for multiple comparisons.

**Figure 2:**
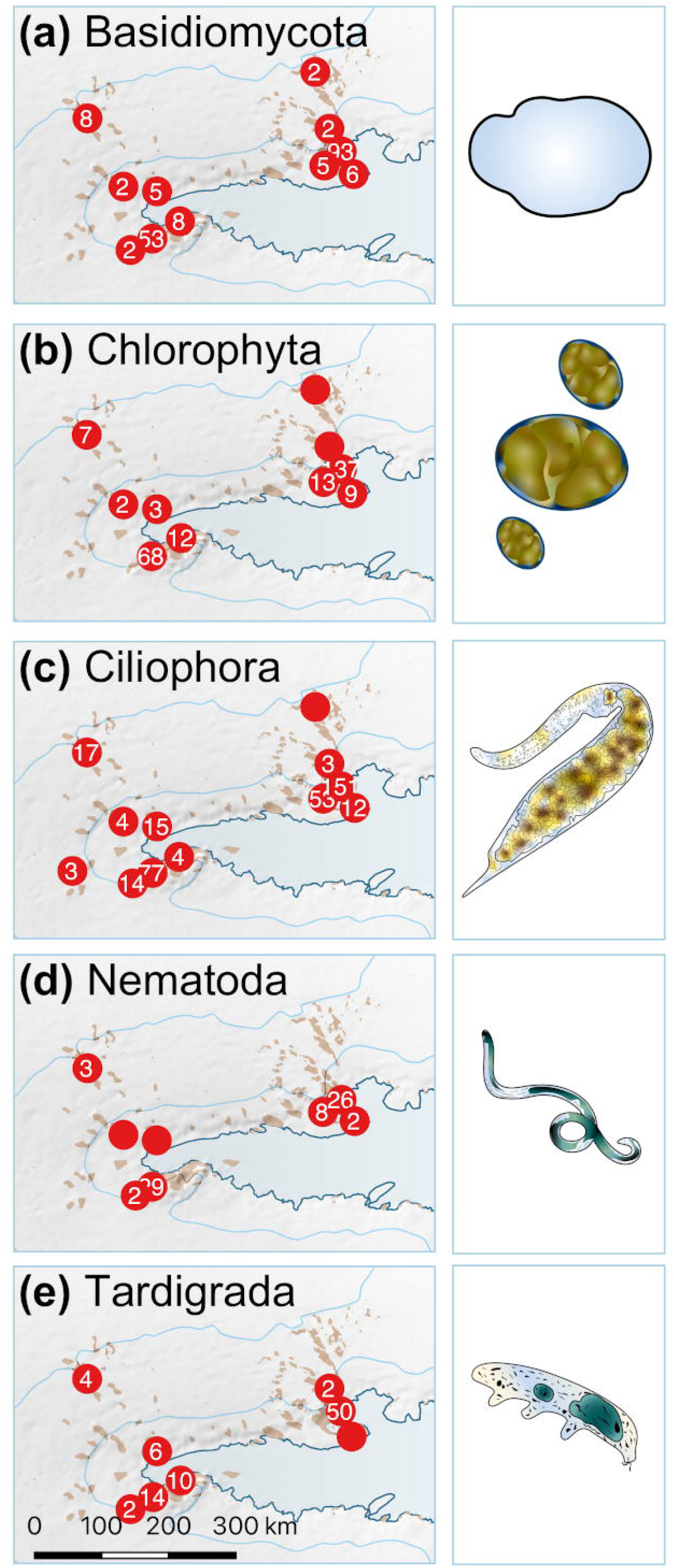
Counts of amplicon sequence variants for phyla deemed significantly influenced by substrate composition (left) and examples of taxonomic assignments (right). The employed relatively short primer pair resulted in survey data encompassing diverse soil life forms of various phyla, at the expense of low-level taxonomic certainty, see Web Table 4 for alignment qualities. (a) *Mrakia frigida* (Basidiomycota; prefect alignment) is closely related to a recently described Antarctic species (Xin and Zhou 2007). (b) *Chloroidium angustoellipsoideum* (Chlorophyta; perfect alignment) is in the same genus as the recently described *Chloroidium antarcticum* (Darienko *et al.* 2018). (c) For *Dileptus jonesi* (Ciliophora; 97.6% identity) possible Antarctic distribution could not be confirmed. Both (d) *Scottnema lindsayae* (Nematoda; perfect alignment) and (e) *Mesobiotus furciger* (Tardigrada; perfect alignment) are known Antarctic species with good reference data coverage (Velasco-Castrillón *et al.* 2014a). Base layers courtesy of the SCAR Antarctic Digital Database, © 1993–2015 Scientific Committee on Antarctic Research; The National Snow and Ice Data Centre, University of Colorado, Boulder; NASA, Visible Earth Team, http://visibleearth.nasa.gov/; Australian Antarctic Division, © Commonwealth of Australia 2006.

Furthermore, we explored the global distribution of the obtained putative species level assignments among phyla significantly influenced by environmental predictors (see below) by querying BISON (bison.usgs.gov), GBIF (www.gbif.org) and iNaturalist (www.inaturalist.org; see Web Text 1 for detailed methods) with R package *spocc*.

## Results

Keeping in mind the coarse raster resolution and model-like character of the climate data, annual mean climate at MM was coldest (−32 ± 0.3 °C), windiest (10.2 ± 0.05 ms^−1^) and with an intermediate amount of precipitation (86 ± 1 mm), when compared to the other two locations (Web Figure 1). ME exhibited the least amount of precipitation (55.3 ± 7 mm), comparatively low wind speeds (5.4 ± 0.5 ms^−1^), and slightly higher temperatures than MM (−28.4 ± 0.6 mm). Closest to the coast, and exposed, LT appeared influenced by the highest precipitation (136 ± 16 mm), variable but moderate wind speeds (5.5 ± 1.7 ms^−1^) and the highest temperature in the surveyed area (−24.1 ± 1.6 °C). We found our chosen climatic variables strongly correlated with the sampling locations, and to improve predictive power excluded the former from further considerations. Instead, we interpreted the statistical effect of location (below) to be a function of annual mean climatic variables.

Retention of eukaryotes in field-derived samples after filtering yielded 2,285,773 reads across 145 samples, derived from 16,524,031 unfiltered sequences (Web Table 3). Per-sample mean coverage was 9,450 reads (min: 2, median: 2,379, max: 86,804). ASV mean coverage after filtering was 2,984 reads (min: 2, median: 132, max: 207,718; Web Figure 6). Collectively after filtering, 766 ASVs were assigned to 495 species across 25 phyla (Web Table 4). Most prevalent phyla (and among those: most prevalent species) by coverage were Ascomycota *(Acanthothecis fontana)*, Chlorophyta *(Coccomyxa* sp.), Basidiomycota *(Mrakia frigida)*, Ciliophora *(Pseudochilodonopsis quadrivacuolata)*, Nematoda *(Scottnema lindsayae)*, Rotifera *(Embata laticeps)*, and Tardigrada *(Mesobiotus furciger)*. (All taxonomic assignments listed here aligned with reference data without gaps at full coverage, and a bit score of 154.6, apart from bit score of 145.6 for *P. quadrivacuolata*)

We found the distribution of five phyla (26 classes, 59 orders, 100 families, 173 species) across the PCMs to be significantly correlated with the considered soil predictors (Figure 2, Web Tables 5 and 6, Web Figure 7). Those taxa were defined by 265 ASVs across 1,210,855 sequences and 142 samples (23 MM, 64 ME, 55 LT). Per-sample mean coverage was 9,460 (min: 2, med: 3863, max: 84,892), per-ASV mean coverage was 4,596, (min: 2, median: 157, max: 128,358; Web Figure 6).

For each predictor significantly correlating with a phylum’s presence (Web Figure 8) we report the expected effect on phylum presence corresponding to one standard deviation (σ) increase of the predictor from its mean (μ), with all other variables held at mean μ. Key significant results included:

i. Low levels of Basidiomycota (62 putative species assignments, Figure 2a) in high pH environments (μ = 7.15, σ = 0.88, E[present _μ_] = 0.6 and E[present _μ +1σ_] = 0.4), and a strong positive relationship of this phylum with dolomite (μ = 0.025 %, σ = 0.05 %, E[present _μ +1σ_] = 0.7).
ii. Very low levels of Chlorophytes (47 species, Figure 2b) at MM plausibly attributable to harsh environmental conditions encountered there (see Supplemental Materials; E[present _LT_] = 0.61 and E[present _MM_] = 0.32, including more alkaline substrates (E[present _μ +1σ_] = 0.46)
iii. Very low levels of Ciliophorans (47 species, Figure 2c) at MM (E[present _LT_] = 0.70 and E[present _MM_] = 0.39), in Sulphur-rich substrates (μ = 528 mg kg ^−1^, σ = 1410 mg kg ^−1^, E[present _μ +1σ_] = 0.61), and in areas relatively rich in pyroxene, amphibole or garnet (μ = 4 %, σ = 4 %, E[present _μ +1σ_] = 0.52)
iv. Very low levels of nematodes (8 species, Figure 2d) at MM (E[present _LT_] = 0.47 and E[present _MM_] = 0.28), and in highly conductive substrates (μ = 0.55 dSm^−1^, σ = 1.07 dSm^−1^, E[present _μ +1σ_] = 0.35)
v. Very low levels of tardigrades (9 species, Figure 2e) in alkaline substrates (E[present _μ_] = 0.22, E[present _μ +1σ_] = 0.14)

Observed fractions of non-zero coefficients are shown Table 1 and Web Figure 8. (95% non parametric bootstrap confidence intervals for non-0 estimates also provided in Web Figure 8.) Directions of all predictor effects on all analyzed taxa presences, including insignificant effects, are listed in Web Table 7.

**Table 1:** Numerical summary of significant coefficient estimates for each phylum as obtained through lasso logistic regression.

| phylum | predictor | 95% CI Coefficient |  | 95% CI Odds ratio |  | Proportion<br>of<br>bootstrap<br>replicates<br>not zero |
| --- | --- | --- | --- | --- | --- | --- |
|  |  | lower | upper | lower | upper |  |
| Basidiomycota | Dolomite | 0 | 1.32 | 1 | -3.70 | 0.93 |
|  | PH | -1.54 | 0.46 | 0.21 | -0.63 | 1.00 |
| Chlorophytes | MM* | -1.32 | -0.10 | 0.26 | 0.90 | 0.99 |
|  | PH | -1.28 | -0.10 | 0.28 | 0.91 | 0.99 |
| Ciliophora | Garnets | -2.07 | -0.11 | 0.13 | 0.90 | 0.99 |
|  | MM | -1.22 | 0.00 | 0.29 | 1.00 | 0.93 |
|  | Sulphur | -3.14 | 0.00 | 0.04 | 1.00 | 0.85 |
| Nematodes | Cond | -2.17 | 0.00 | 0.11 | 1.00 | 0.99 |
|  | MM | -2.10 | -0.26 | 0.12 | 0.77 | 0.99 |
| Tardigrades | PH | -1.42 | 0.00 | 0.23 | 1.00 | 0.95 |

For 66 of our 173 putative species assignments 778 georeferenced records could be obtained (of those 65% from GBIF, 27% iNaturalist, 7% BISON). Of the obtained 123 locations 4% were in Africa, 1.6% in Antarctica, 13% Asia, 32% Europe, 21% North America and 10% in South America (Web Figures 9 and 10, Web Table 7). The sole species recorded for Antarctica (here: south of 66.56°) was the nematode *Scottnema lindsayae*. Observations north of the polar circle (likewise 66.56°) included Basidiomycota *(Gloiocephala aquatica, Stereum rugosum, Mrakia frigida, Rhodotorula mucilaginosa)*, Chlorophyta *(Haematococcus lacustris, Oophila amblystomatis)*, and Ciliophora *(Furgasonia blochmanni, Chilodonella acuta (Ciliophora), Tachysoma pellionellum)*. Refer to Web Tables 4 and 5 for alignment qualities.

## Discussion

Our Antarctic case study demonstrates two key technologies to be useful for baseline biodiversity surveys across large spatial scales in extremely remote environments – robust predictive statistics, such as the Lasso, now often used in machine learning algorithms (Muthukrishnan and Rohini 2016), as well as biodiversity information derived from environmental DNA (Czechowski *et al.* 2017). To the best of our knowledge, our work is the first in associating environmental DNA data to environmental predictors by means of the Lasso to yield accurate detection probabilities for taxonomic groups, also in Antarctica. Thus, we present an analytical framework to identify areas for targeted species-level biodiversity surveys, using other markers, or predictors for Antarctica, and possible other hardly accessible locations.

Our expanded analyses of the original raw data (Czechowski *et al.* 2016b) made use of new algorithms for processing environmental DNA sequences (eg Callahan *et al.* 2016, 2017), along with more extensive reference databases for taxonomic assignment, and new algorithms available with R (R Core Development Team 2019). While our results are in line with earlier findings relating eukaryote distribution to their environment in the PCMs and Antarctica (eg Czechowski *et al.* 2016a, b; Bottos *et al.* 2020), our approach adds accuracy to those findings with respect to five phyla.

A strength of our analyses is the relatively easy retrieval of biological survey data encompassing many phyla (probably including many cryptic and unknown species) across many samples. The weakness of the employed 18S marker is its limited ability to discern many distinct sequence variants on a low taxonomic (eg species) level. Regardless, identification of species with likely Antarctic occurrence such as the known Antarctic nematode *Scottnema lindsayae* and tardigrade *Mesobiotus furciger* by means of a relatively short and highly conserved primer pair highlights the ability of environmental DNA to retrieve species occurrence records, provided that sufficient sequence data is available for taxonomic assignment. Consequently, we believe that environmental DNA analysis should be the method of choice to obtain biodiversity data from Antarctica, particularly when many samples are to be analyzed, but other markers are needed to investigate fine scaled endemism, and to obtain better taxonomic resolution.

Georeferencing our putative species assignments by means of publicly accessible databases had limited success. The limitations of reference databases became obvious when known Antarctic species, such as *Acutuncus antarcticus* (Web text 1), the latter identified among our data through a perfect alignment with bit score 154.6, were not found, and only 38% of all putative Antarctic species assigned by us were georeferenced at all. High occurrence prevalence in North America, and Europe indicates sampling bias in GBIF, iNaturalist and BISON and highlight a substantial weaknesses of publicly accessible global biodiversity data concerning cryptic eukaryotes.

Eukaryotic distribution patterns reported in related Antarctic studies provide context for our observations from the PCMs. The rarity of Chlorophytes, Ciliophorans, and the otherwise ubiquitous nematodes at MM in relation to the two other lower altitude and more northerly locations (ME, LT) seem to confirm trends of increasing eukaryotic richness and diversity with decreasing latitude and altitude (Czechowski *et al.* 2016a; Thompson *et al.* 2020; Zhang *et al.* 2020), but such patterns are not always evident at the scales investigated here. Rather, Antarctic biodiversity can be surprisingly regionalized (Convey *et al.* 2014) and correspondingly, our study finds surprisingly high eukaryotic diversity to unexpectedly occur even in the harshest environments, such as local ice-soil substrate boundaries at Mount Menzies (Figures 1a, 2). The absence of ciliophorans from Sulphur-rich substrates, and of nematodes from highly conductive soil interstices matches findings of distribution patterns being shaped by age-related salt accumulation at the surface-air interface of frozen soils described with other analytical approaches (Velasco-Castrillón *et al.* 2014b; Lee *et al.* 2019).

In absence of other predictors, our study highlights the importance of neutral substrate pH, low conductivity, and key minerals (dolomite, pyroxene, amphibole, or garnet) to predict high eukaryote density in Antarctic substrates. We corroborate the negative influence of substrate alkalinity on Antarctic Basidiomycota (Arenz and Blanchette 2011). Bioregionalization notwithstanding, distance to coast once more appears as suitable proxy variable negatively related to the presence of chlorophytes and ciliophorans (Thompson *et al.* 2020). Additionally, we find soil alkalinity, Sulphur content and substrates pyroxene, amphibole, or garnets to constrain distribution of the former. Among nematodes, our results (i.e. perfect alignment between our Antarctic 18S sequence from Mount Menzies and an annotated reference sequence) indicate that *Scottnema lindsayae* could likely occur in high altitude and high latitude environments such as MM, but then would be influenced by the species’ general indifference (rather than affinity, compare Zawierucha *et al.* 2019) to alkaline substrates, and must be highly localized (at least at MM) if encountered at high abundance (Smykla *et al.* 2018; Zawierucha *et al.* 2019). Lastly, we confirm the negative association between tardigrade occurrence and alkaline substrates observed in Victoria Land (eg Smykla *et al.* 2018).

Based on our findings, ice-free areas with high annual mean precipitation, low wind speeds and relatively high temperatures, exhibiting substrates with a neutral pH and low conductivity, which are rich in dolomite but poor in pyroxene, amphibole, or garnets, are likely to be highly biodiverse in the Antarctic and should harbor candidates for more focused conservation management and higher resolution DNA markers with morphological species level investigations. Furthermore, locations with more extreme environmental conditions may harbor endemic relic fauna equally warranting protection (Convey *et al.* 2014). Our results are in line with observations in other (including polar and alpine) ecosystems, where soil pH was found to be an important factor determining bacterial and fungal community (Siciliano *et al.* 2014; Bottos *et al.* 2020). At the same time, Antarctic soil ecosystems are relatively simple and are assumed to mostly lack complex biotic interactions, although such interactions may be more present in coastal terrestrial ecosystems (Velasco-Castrillón *et al.* 2014b; Lee *et al.* 2019). Consequently, the soil eukaryote distribution patterns observed especially at Mount Menzies are likely predominantly shaped by abiotic factors and would be gradually more influenced by limited biotic interactions, lower latitude substrates or more costal substrates (ME, LT).

## Conclusion

We provide a case study highlighting the utility of environmental molecular data and predictive analysis algorithms to inform on the presence of eukaryote taxa by means of relatively easily measured soil predictors, which can be combined with readily available climate data. Rather than recognizing trends, our analytical technique provides accurate detection probabilities for Basidiomycota, Chlorophytes, nematodes, and tardigrades in relation to bedrock mineral composition, pH, conductivity, Sulphur contents, and arguably, overall harshness of environmental conditions. These, here quantified, relationships enable more precise distribution modeling of phylum presences over large spatial scales. Our approach may be used identify regions worthy of species level biodiversity surveys, possibly employing faster evolving molecular markers or logistically more challenging morphologic biodiversity assessments. We believe our approach to be valuable to inform further development and understanding of both Antarctic biogeography and conservation areas.

## Supporting information

Web Figure 1

Web Figure 2

Web Figure 3

Web Figure 4

Web Figure 5

Web Figure 6

Web Figure 7

Web Figure 8

Web Figure 9

Web Figure 10

Web Table 1

Web Table 2

Web Table 3

Web Table 4

Web Table 5

Web Table 6

Web Table 7

Web Table 8

Web Text 1

## Acknowledgments

P.C. was supported by The University of Adelaide through an International Post-Graduate Research Scholarship, through the Royal Society of South Australia, and the University of Otago. M.S. received AAS funding from The Australian Antarctic Division, science project 2355 and A.T. was supported by AAS 4296. Alan Cooper and M.S. received funding for this project through Australian Research Council linkage grant LP0991985. M.S. and P.C. received funding for this project from the Sir Mark Mitchell Foundation and the University of Otago. M.K was funded by the University of Otago. We thank Robert McPhee for the artistic contribution of Figure 2. We thank Diana Wall for providing helpful comments on an earlier draft of the manuscript. Authors’ contributions to this article conform with the CASRAI Contributor Roles Taxonomy (https://casrai.org/credit).

