## Supplementary material for "Antarctic biodiversity predictions through substrate qualities and environmental DNA": Web Figure 1

**Web Figure 1:** Upper: Climatic variables (van Wessem *et al.* 2014; Van Wessem *et al.* 2014; Matsuoka *et al.* 2021) in study extent after bilinear desegregation to 1 km pixel size. Indicated are sampling points and 20 km buffers for variable extraction. Lower: Extracted climatic variables per location, the median value in the buffered location was used.

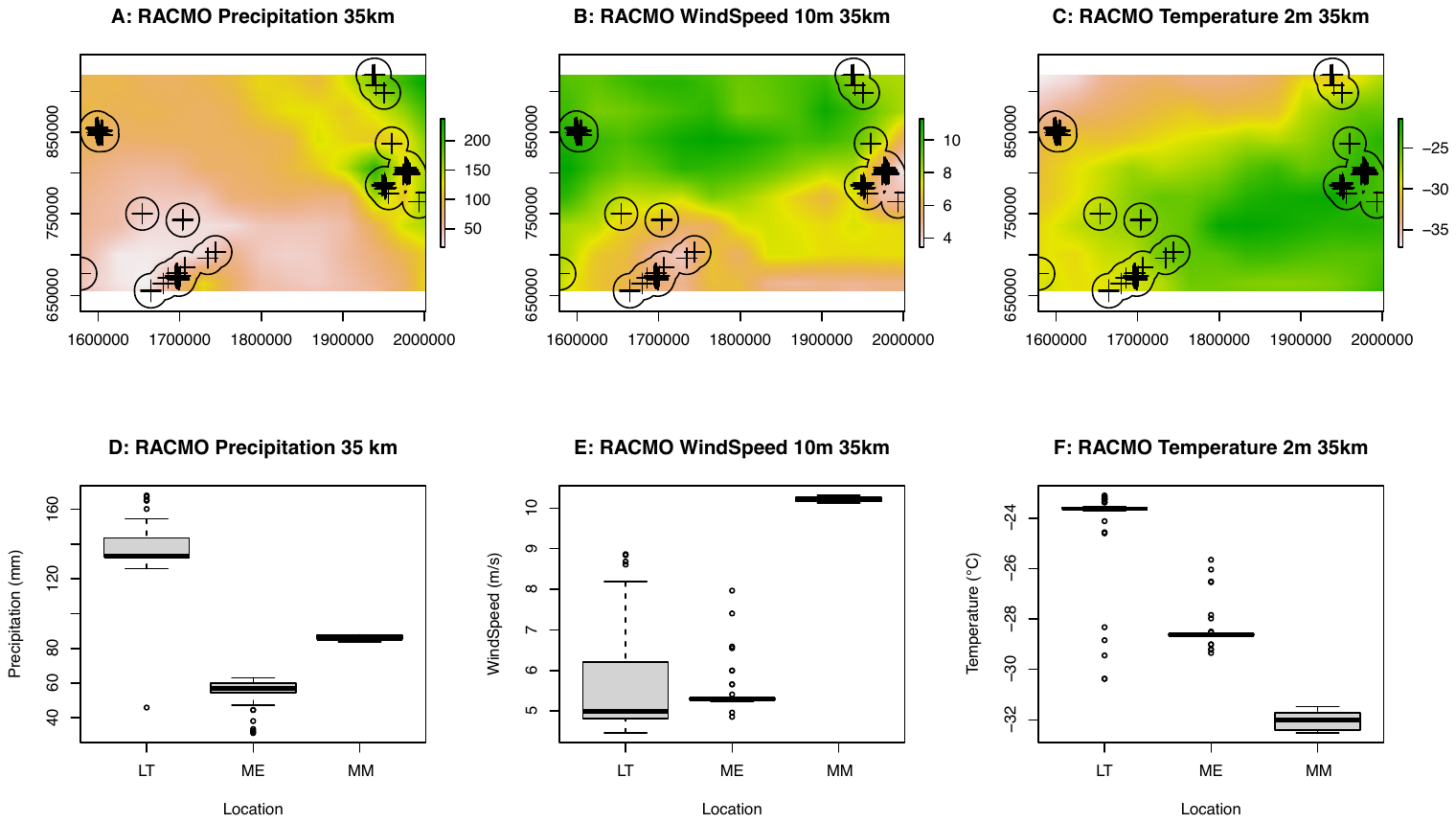
