## Supplementary material for "Antarctic biodiversity predictions through substrate qualities and environmental DNA": Web Figure 2

**Web Figure 2:** Density of raw predictors preceding data filtering. Also shown are predictors that were excluded due to missing data. Count of successful observations in parentheses, and further detailed in WebTable 7. “AGE KA”: bedrock age in ka, estimated with geological maps; “AMMN” Ammonium N content (mg/Kg); “CALCITE”: calcite fraction % as integrated from X-ray diffraction spectra; “CARB”: organic carbon in %; “CHLORITE”: chlorite % as integrated from X-ray diffraction spectra; “COND”: conductivity in dS/m; “DOLOMITE”: Dolomite fraction % as integrated from X-ray diffraction spectra; “ELEVATION”: as measured with handheld GPS on site; “GARNET”: pyroxene, amphibole or garnet fraction % as integrated from X-ray diffraction spectra; “KAOCHLOR”: kaolin, chlorite and chlorite fraction % as integrated from X-ray diffraction spectra; “LatDEC and “LongDEC”, latitude and longitude in decimal degrees as measured with handheld GPS (WGS84) on site; “MICAS”: micas fraction % as integrated from X-ray diffraction spectra; “NITR”: Nitrate N (mg/Kg); “PH CACL” and “PH H2H”: pH values as measured for calcium dichloride and water; “PHOS” and “POTA”: phosphorus and potassium (mg/kg); “QUARTZ”: quartz % as integrated from X-ray diffraction spectra; “RACMO precip mm 35to1km”, “RACMO windsp 10m 35to1km” and “RACMO tmp 2m 35to1km”: precipitation, windspeed and temperature as extracted from climate rasters (also see Fig. 3); “RibLibConcAvg”: DNA concentration of pooled library replicates prior to equimolarisation; “RibPoolConcAvg”: DNA concentration of pooled library replicates after equimolarisation; “RLU”: relative light units across two replicates, as measured with an ATP meter during field work; “SLOPE”: Slope angle as measured at sampling site. “SULPH”: Sulphur content (mg/kg); “TEXT” coarseness of substrate. “TTITANITE”: titanite fraction % as integrated from X-ray diffraction spectra.

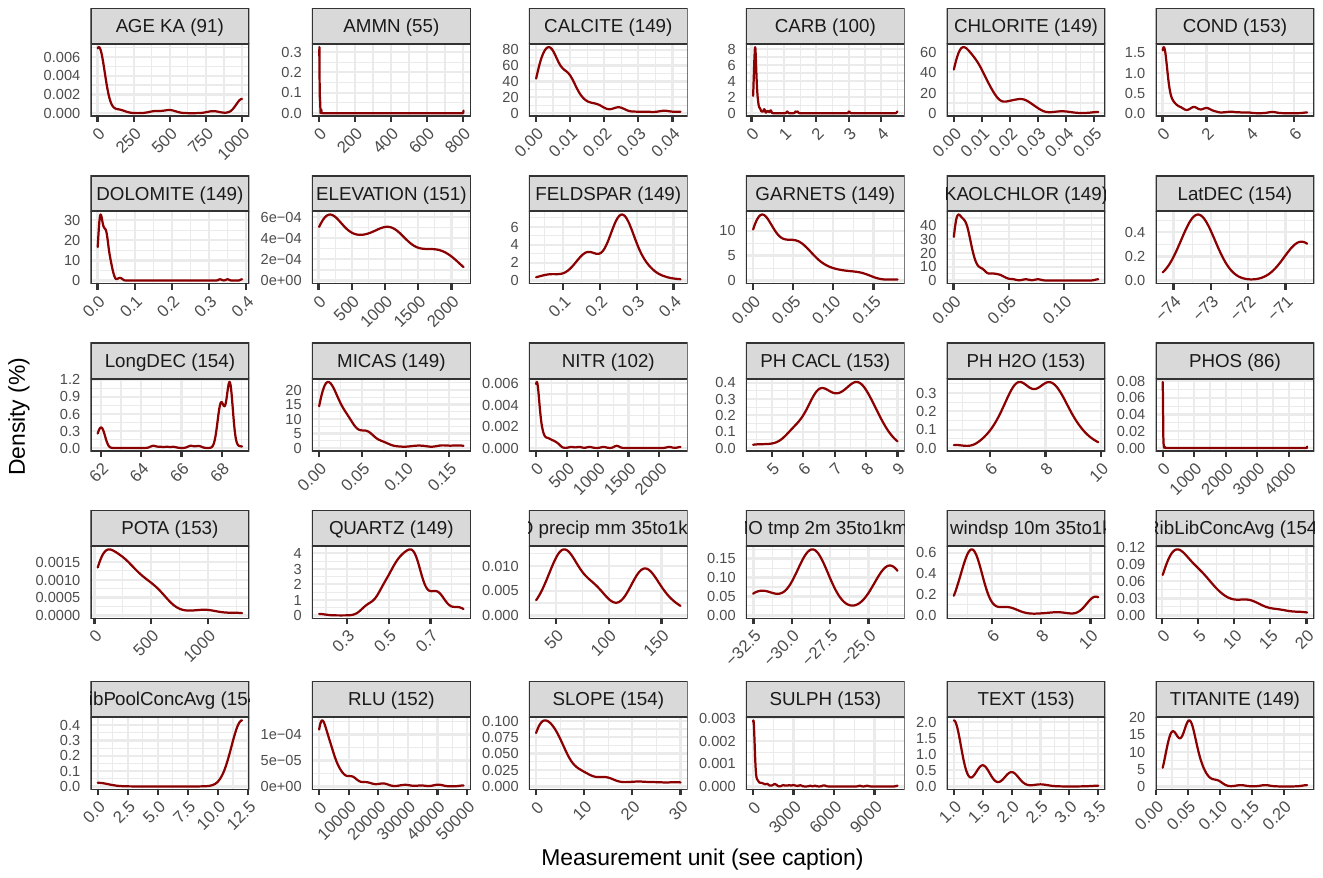
