## Supplementary material for "Antarctic biodiversity predictions through substrate qualities and environmental DNA": Web Figure 3

**Web Figure 3:** Denoising and filtering statistics to obtain raw data as provided by Qiime (Bolyen *et al.* 2019)

**Sub-Figure a:** first plate

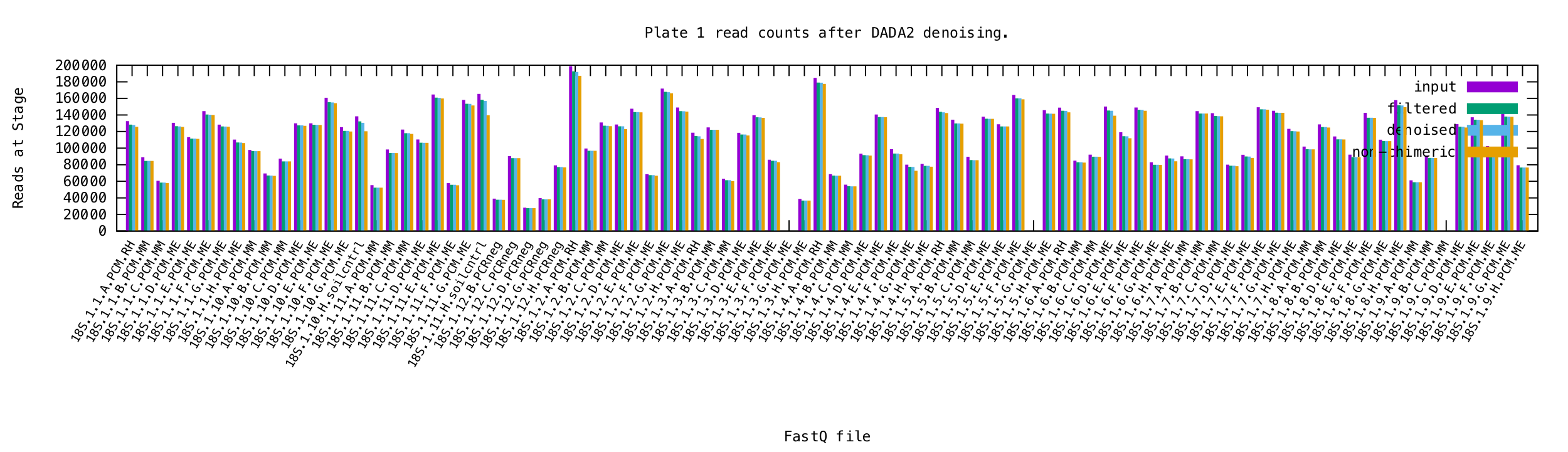

**Sub-Figure b:** second plate

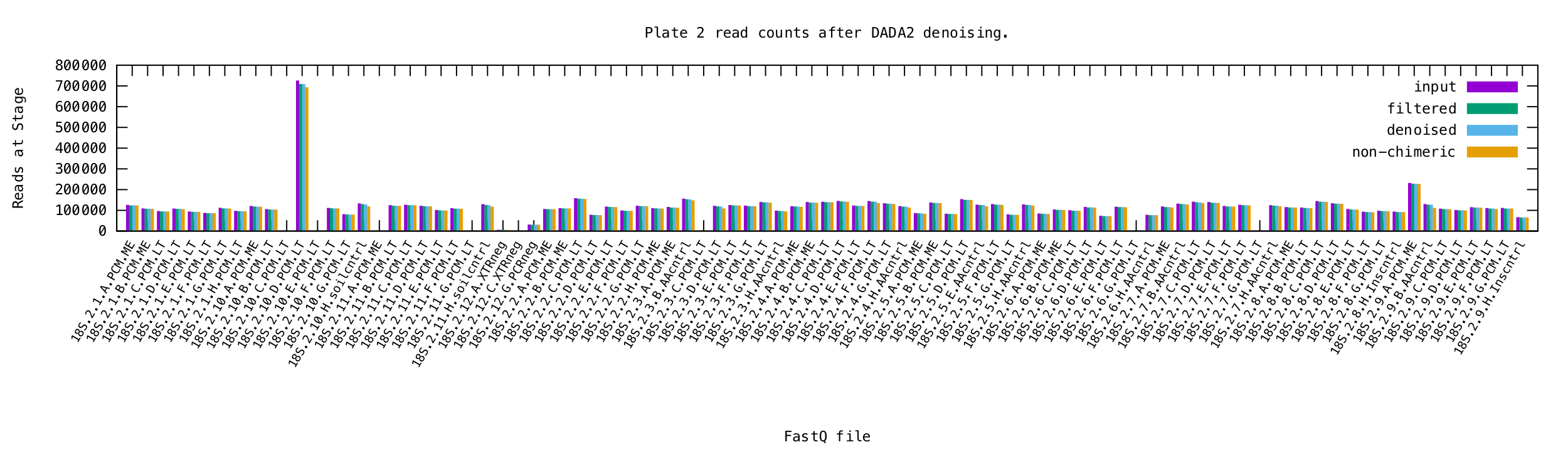
