## Supplementary material for "Antarctic biodiversity predictions through substrate qualities and environmental DNA": Web Figure 4

**Web Figure 4:** Taxonomic composition overview of environmental Amplicon Sequence Variants before and after filtering stages. All species and reads contained in controls where removed from remaining data prior to further analysis.

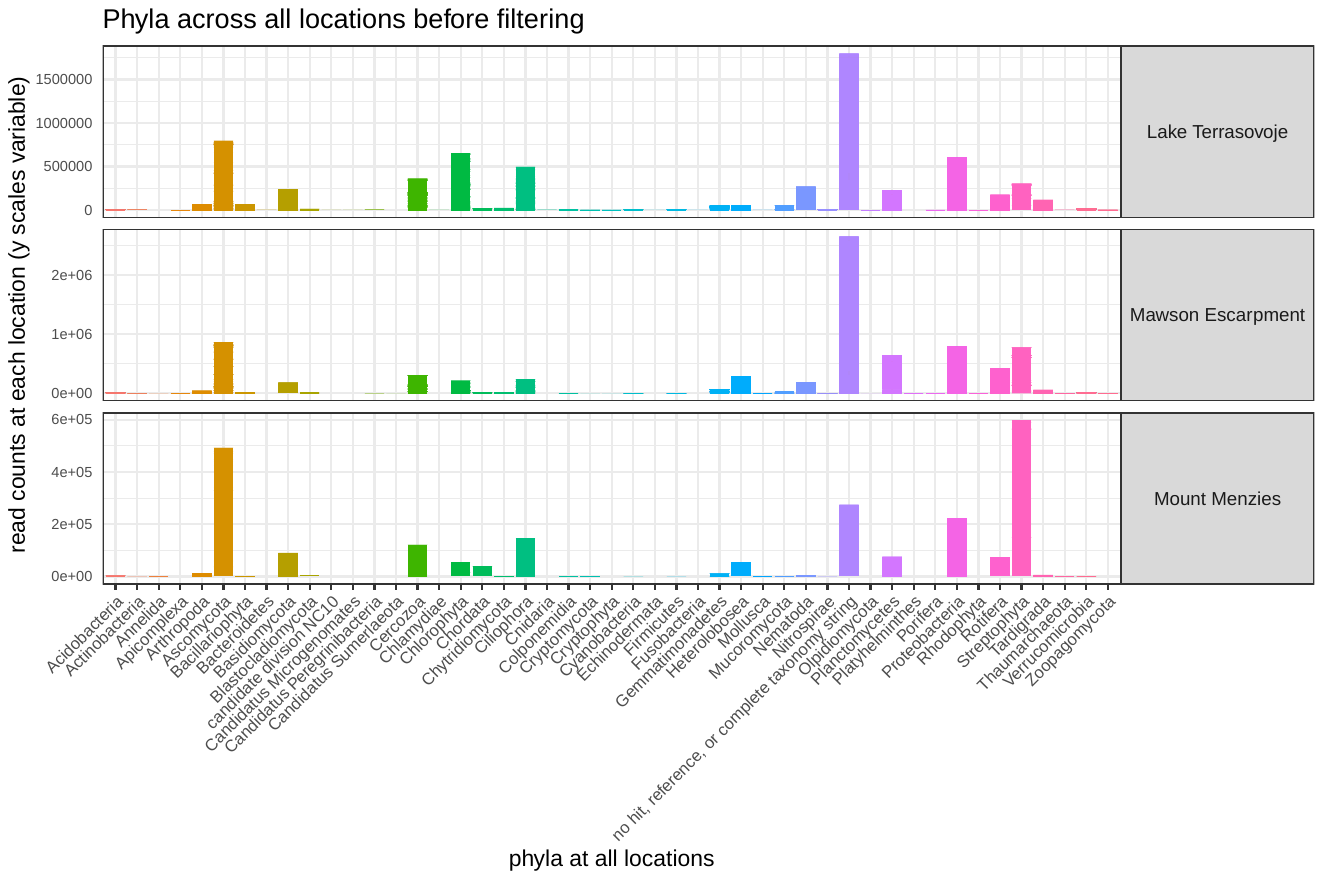

**Sub-Figure a:** Prior to filtering

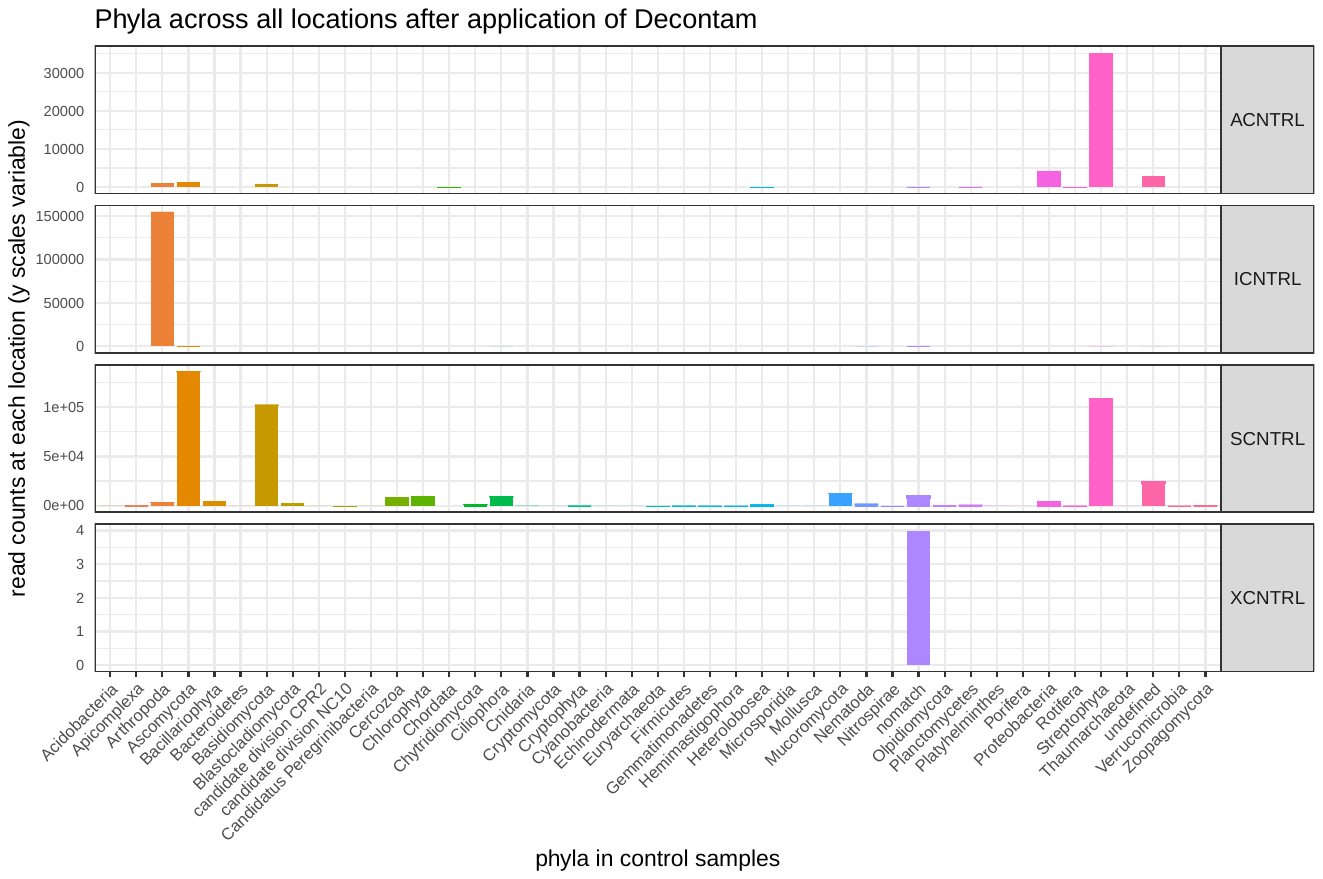

**Sub-Figure b:** Sequences remaining in control reaction after application of decontamination using R package *decontam* (Davis *et al.* 2018). ACNTROL: Amplification control; ICNTRL: non-Antarctic insect positive control; SCNTRL: Australian soil positive control; XCNTRL: Extraction control (Czechowski *et al.* 2016a, b)

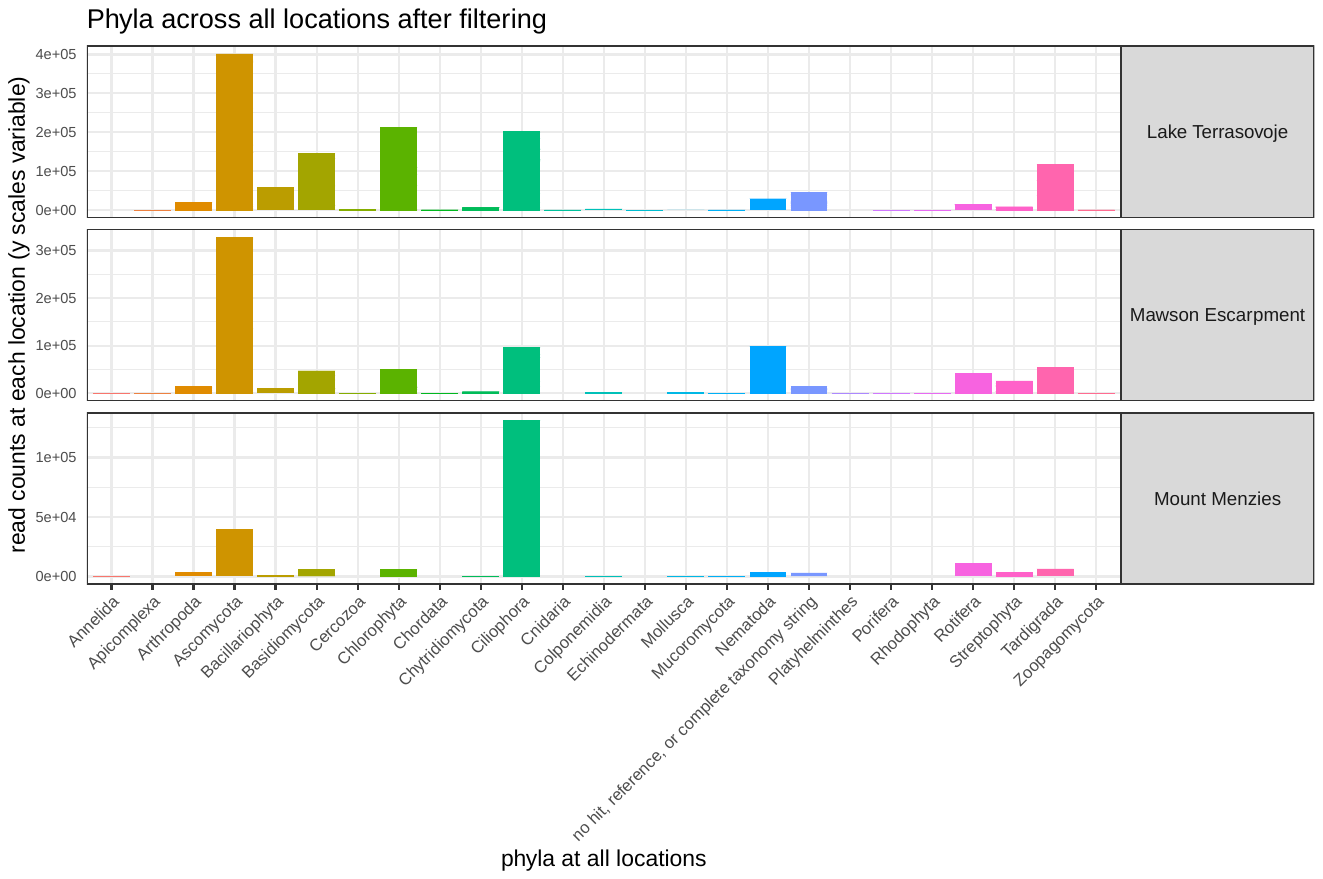

**Sub-Figure c:** Taxonomic assignments across 2 285 773 environmental 125 bp 18S Amplicon Sequence Variants (*sensu* Callahan *et al.* 2017) following NCBI hierarchy. Assignments derived from references sequences at least 50% identical to query sequences, assignment certainty e of 10-10, for matches of at least 90% query coverage, excluding environmental sequences. Collectively, ASVs were assigned to 495 species across 25 phyla.
