## Supplementary figures and images for "Antarctic biodiversity predictions through substrate qualities and environmental DNA"

### Web Figure 5

**Web Figure 5:** Predictor overview after data filtering, per location**.**

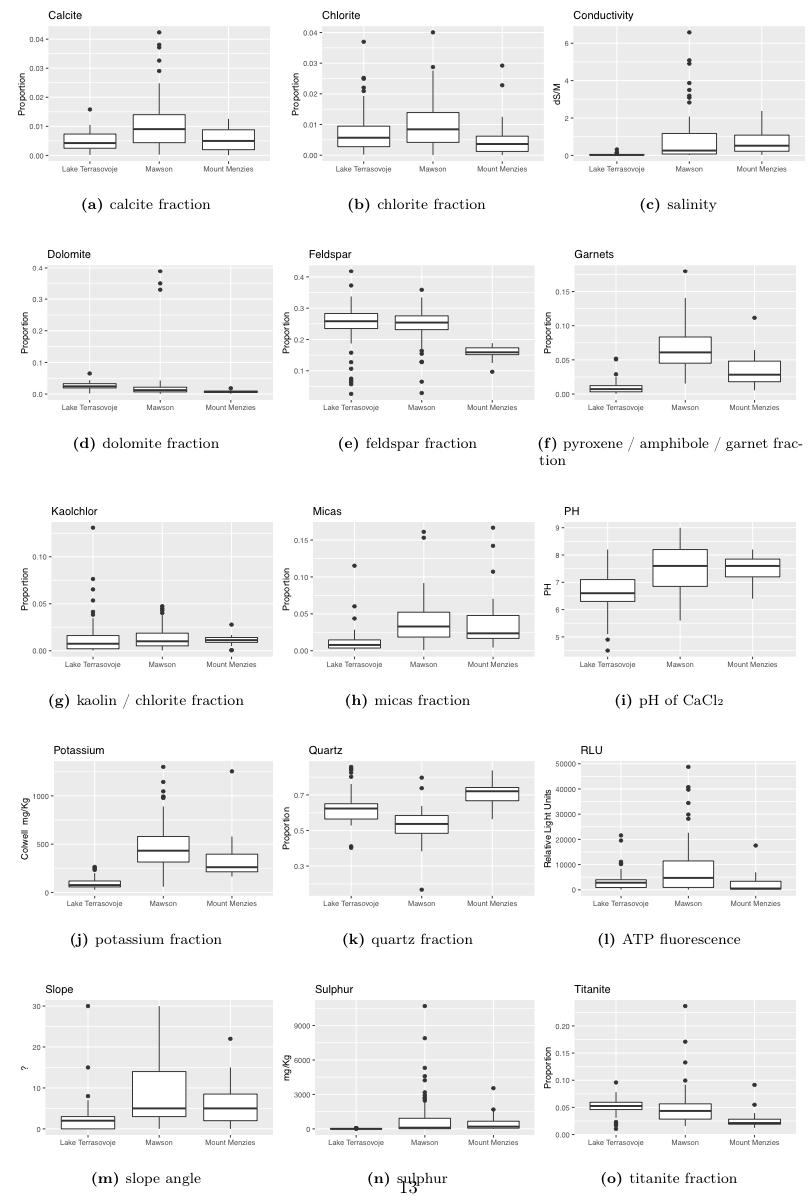

### Web Figure 6

**Web Figure 6:** Sample and ASV coverages prior to and after filtering stages.

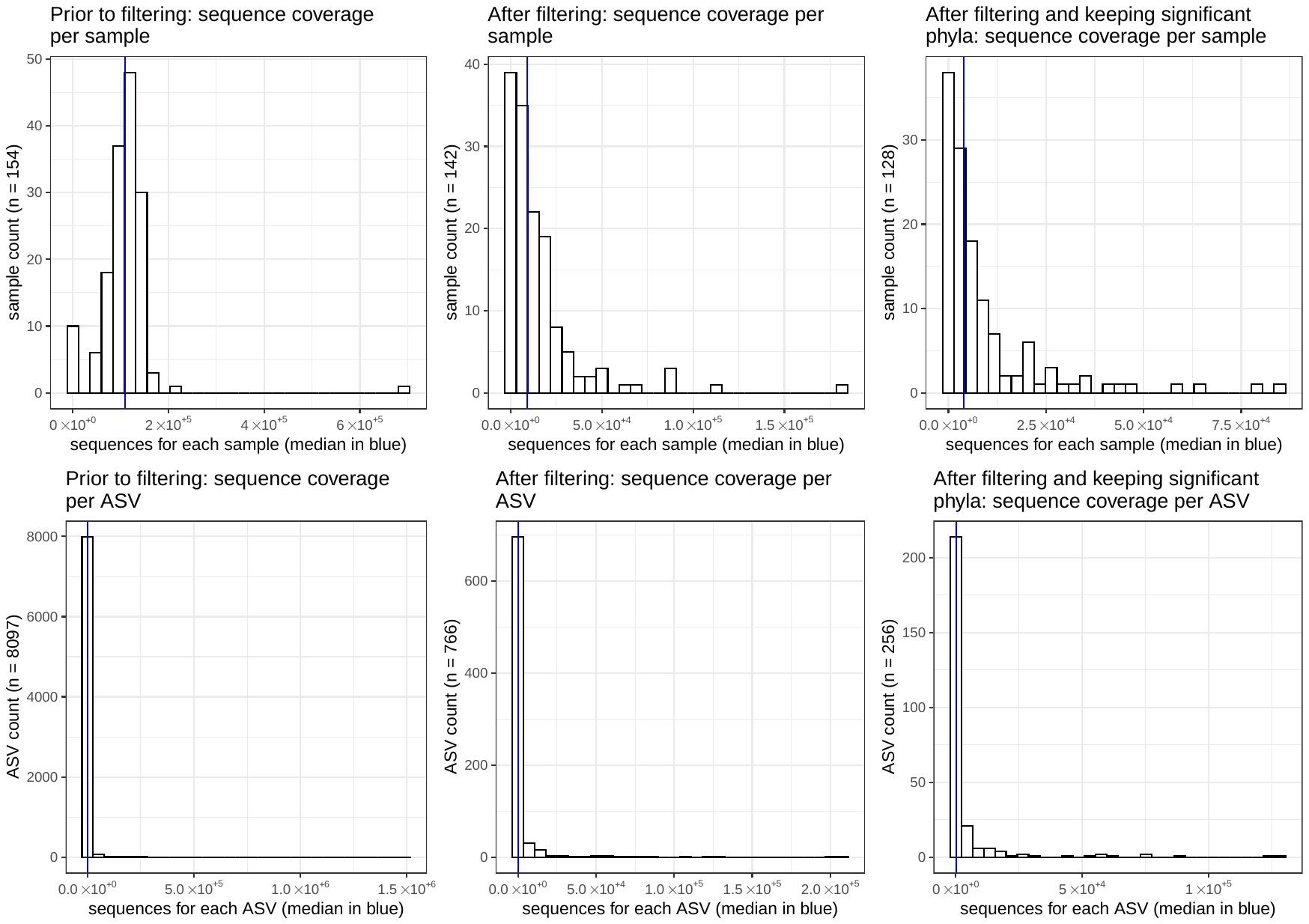

### Web Figure 7

**Web Figure 7:** Full regression results.

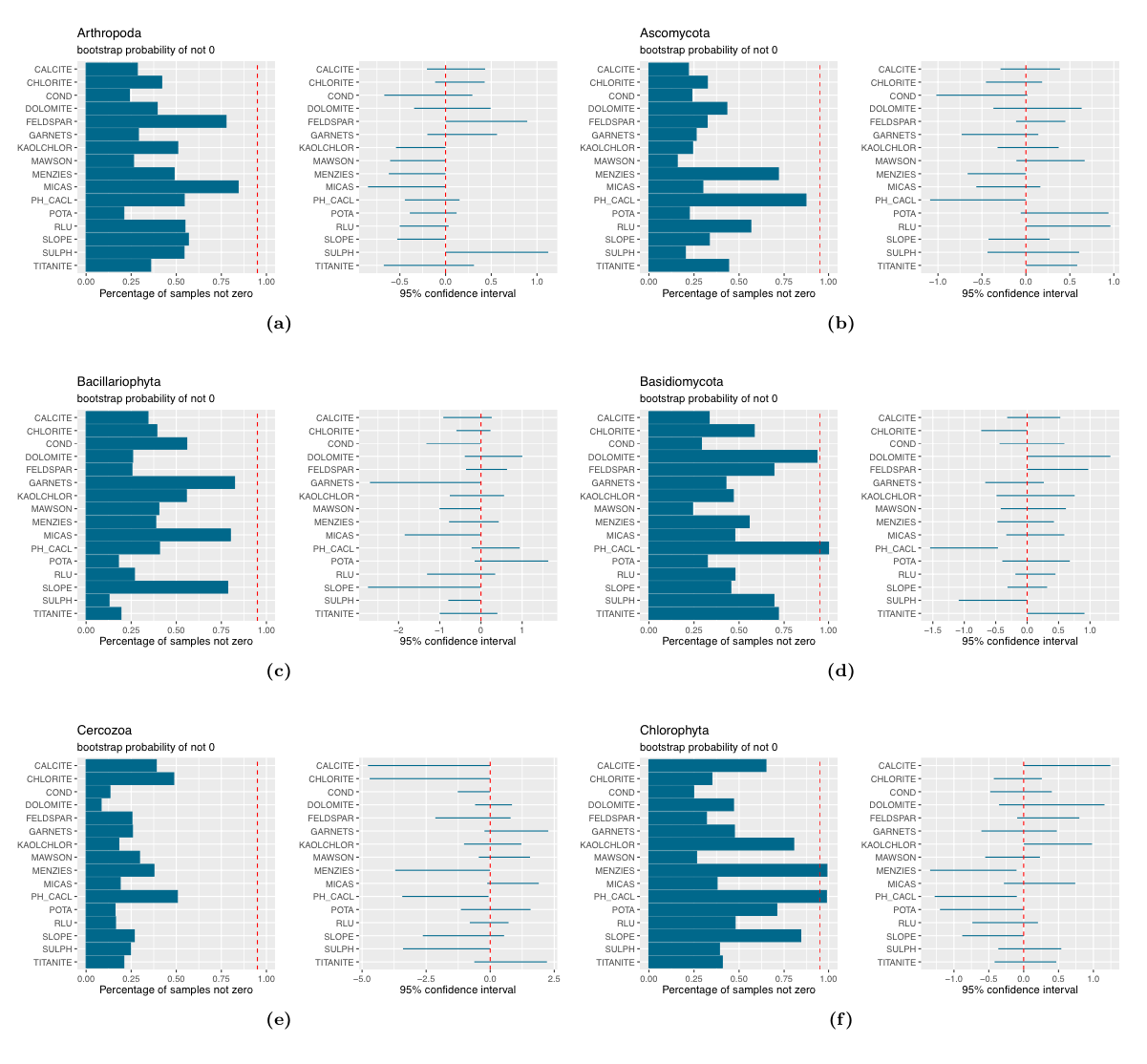

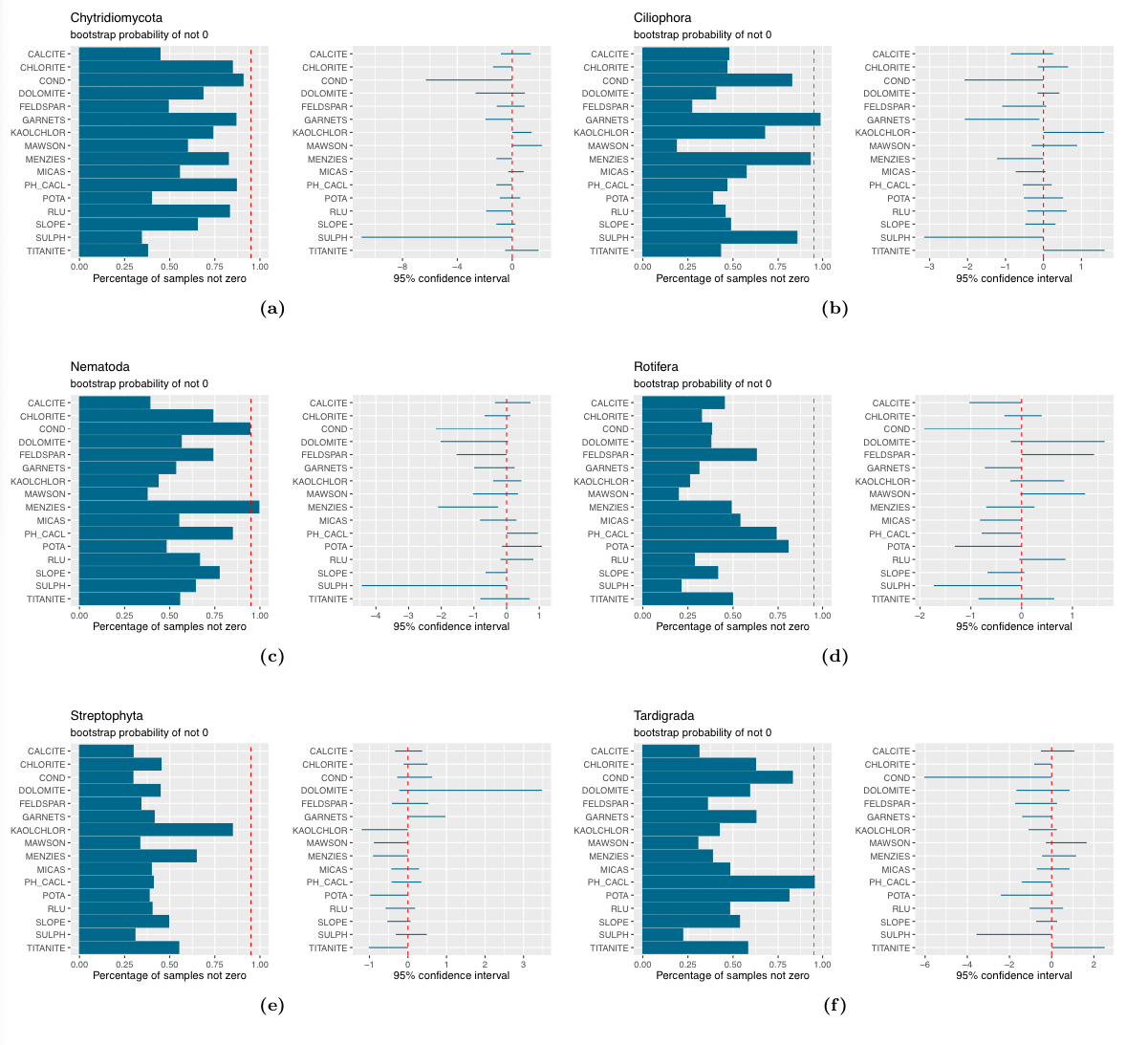
