## Supplementary material for "Antarctic biodiversity predictions through substrate qualities and environmental DNA": Web Figure 8

**Web Figure 8:** Subset of phyla with distributions significantly correlated with analyzed environmental predictors (also compare Table 1 and Supporting Information). Left panels: Proportions of bootstrap (n = 10 000) samples with non- zero estimates and delineation of 95% confidence levels. Right panels: Confidence intervals for estimates (significant predictors should not include 0).

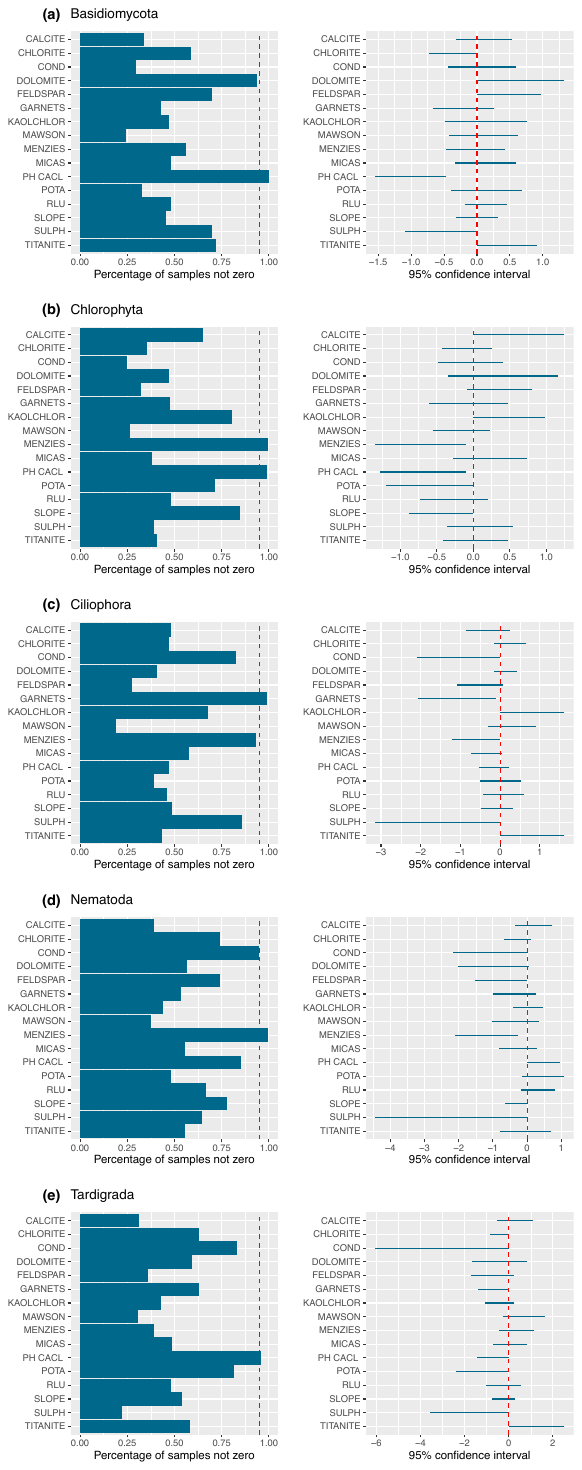
