## Supplementary material for "Antarctic biodiversity predictions through substrate qualities and environmental DNA": Web Figure 9

**Web Figure 9:** Location of 778 occurrence records obtained from BISON (United States Federal Resource for Biological Occurrence Data, <https://bison.usgs.gov>; n = 53), GBIF (Global Biodiversity Information Facility, <https://www.gbif.org>; n = 523), and iNaturalist (<https://www.inaturalist.org>; n = 202) for 66 results of 173 submitted queries, listed by continent of record and occurrence.

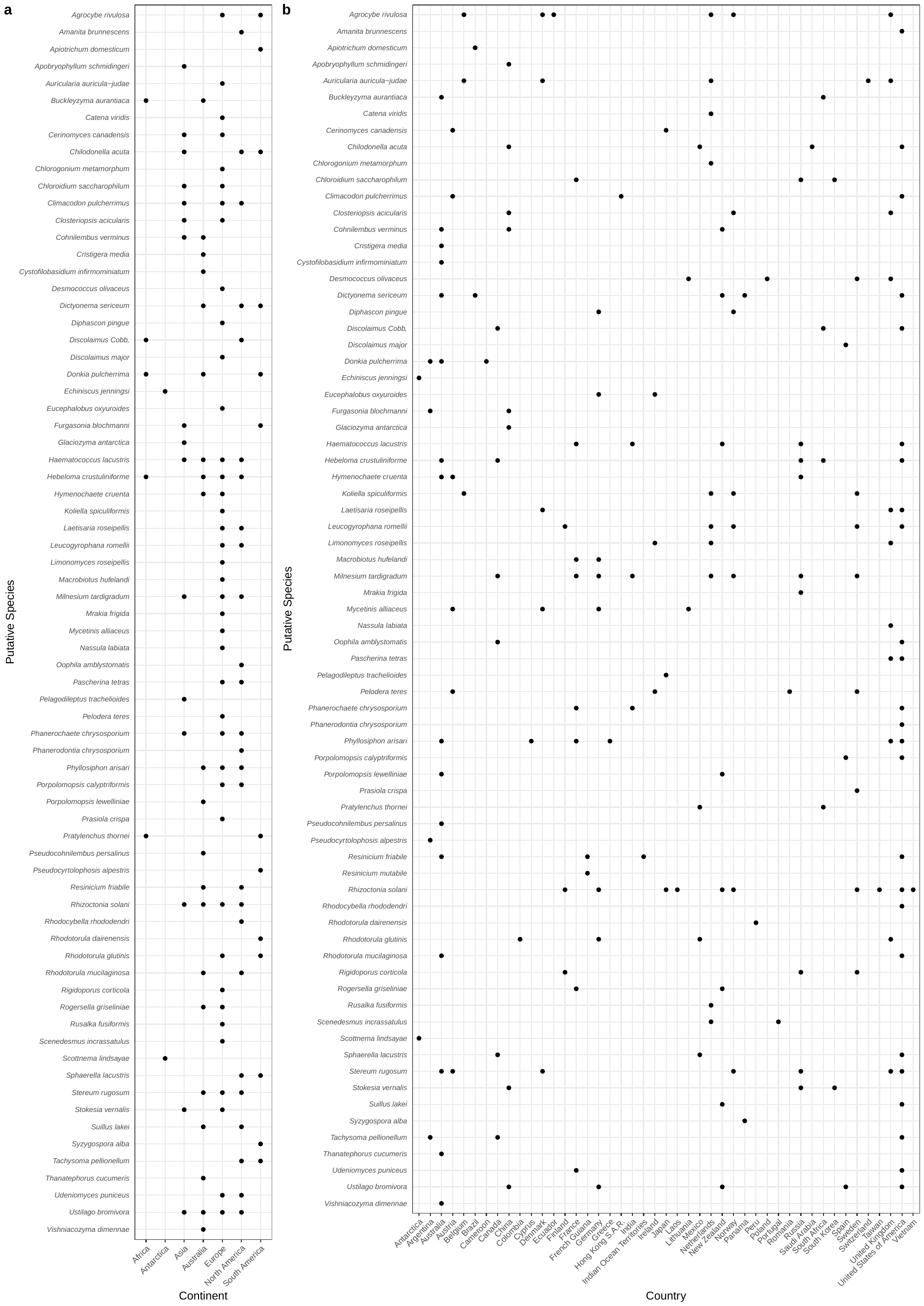
