## Supplementary material for "Antarctic biodiversity predictions through substrate qualities and environmental DNA": Web Table 1

**Web Table 1:** Sequence library names and locations of filtered data, also used to generate Figure 1 in main text.

| Sample | Location | Longitude | Latitude |
| --- | --- | --- | --- |
| 18S.1.1.C.PCM.MM | Mount Menzies | 62,02444444 | -73,38888889 |
| 18S.1.10.A.PCM.MM | Mount Menzies | 61,88722222 | -73,42555556 |
| 18S.1.10.B.PCM.MM | Mount Menzies | 62,01055556 | -73,39277778 |
| 18S.1.10.C.PCM.MM | Mount Menzies | 62,12638889 | -73,43972222 |
| 18S.1.11.A.PCM.MM | Mount Menzies | 61,84666667 | -73,42916667 |
| 18S.1.11.B.PCM.MM | Mount Menzies | 62,15555556 | -73,43222222 |
| 18S.1.11.C.PCM.MM | Mount Menzies | 61,8775 | -73,42027778 |
| 18S.1.2.B.PCM.MM | Mount Menzies | 61,84388889 | -73,41777778 |
| 18S.1.2.C.PCM.MM | Mount Menzies | 62,14972222 | -73,42333333 |
| 18S.1.3.B.PCM.MM | Mount Menzies | 61,84305556 | -73,42333333 |
| 18S.1.3.C.PCM.MM | Mount Menzies | 62,04222222 | -73,3875 |
| 18S.1.4.B.PCM.MM | Mount Menzies | 61,85916667 | -73,43944444 |
| 18S.1.5.B.PCM.MM | Mount Menzies | 61,98027778 | -73,42444444 |
| 18S.1.5.C.PCM.MM | Mount Menzies | 62,09611111 | -73,43722222 |
| 18S.1.6.B.PCM.MM | Mount Menzies | 61,95666667 | -73,42583333 |
| 18S.1.6.C.PCM.MM | Mount Menzies | 62,09055556 | -73,43666667 |
| 18S.1.7.A.PCM.MM | Mount Menzies | 61,87111111 | -73,42277778 |
| 18S.1.7.B.PCM.MM | Mount Menzies | 62,01361111 | -73,45111111 |
| 18S.1.7.C.PCM.MM | Mount Menzies | 62,12305556 | -73,44805556 |
| 18S.1.8.A.PCM.MM | Mount Menzies | 62,02166667 | -73,42888889 |
| 18S.1.8.B.PCM.MM | Mount Menzies | 61,94027778 | -73,42916667 |
| 18S.1.9.A.PCM.MM | Mount Menzies | 62,05361111 | -73,4 |
| 18S.1.9.B.PCM.MM | Mount Menzies | 62,15305556 | -73,44333333 |
| 18S.1.1.D.PCM.ME | Mawson Escarpment | 68,31888889 | -73,32027778 |
| 18S.1.1.F.PCM.ME | Mawson Escarpment | 68,40666667 | -73,30666667 |
| 18S.1.1.G.PCM.ME | Mawson Escarpment | 68,38722222 | -73,31722222 |
| 18S.1.1.H.PCM.ME | Mawson Escarpment | 68,50194444 | -73,30833333 |
| 18S.1.10.D.PCM.ME | Mawson Escarpment | 68,45111111 | -73,3075 |
| 18S.1.10.E.PCM.ME | Mawson Escarpment | 68,39861111 | -73,30416667 |
| 18S.1.10.F.PCM.ME | Mawson Escarpment | 68,41111111 | -73,31916667 |
| 18S.1.10.G.PCM.ME | Mawson Escarpment | 68,50361111 | -73,31055556 |
| 18S.1.11.D.PCM.ME | Mawson Escarpment | 68,44666667 | -73,30916667 |
| 18S.1.11.E.PCM.ME | Mawson Escarpment | 68,43194444 | -73,30777778 |
| 18S.1.11.G.PCM.ME | Mawson Escarpment | 68,35361111 | -73,3275 |
| 18S.1.2.D.PCM.ME | Mawson Escarpment | 68,33555556 | -73,31777778 |
| 18S.1.2.E.PCM.ME | Mawson Escarpment | 68,44972222 | -73,30444444 |
| 18S.1.2.F.PCM.ME | Mawson Escarpment | 68,43194444 | -73,30888889 |
| 18S.1.2.G.PCM.ME | Mawson Escarpment | 68,38111111 | -73,3175 |
| 18S.1.2.H.PCM.ME | Mawson Escarpment | 68,34416667 | -73,3275 |
| 18S.1.3.D.PCM.ME | Mawson Escarpment | 68,32388889 | -73,32 |
| 18S.1.3.E.PCM.ME | Mawson Escarpment | 68,44666667 | -73,3075 |
| 18S.1.3.H.PCM.ME | Mawson Escarpment | 68,35416667 | -73,32833333 |
| 18S.1.4.D.PCM.ME | Mawson Escarpment | 68,33416667 | -73,31888889 |
| 18S.1.4.E.PCM.ME | Mawson Escarpment | 68,44166667 | -73,30166667 |
| 18S.1.4.F.PCM.ME | Mawson Escarpment | 68,46666667 | -73,3025 |
| 18S.1.4.G.PCM.ME | Mawson Escarpment | 68,4375 | -73,32027778 |
| 18S.1.4.H.PCM.ME | Mawson Escarpment | 68,37555556 | -73,33 |
| 18S.1.5.D.PCM.ME | Mawson Escarpment | 68,31638889 | -73,32444444 |
| 18S.1.5.E.PCM.ME | Mawson Escarpment | 68,46055556 | -73,30361111 |
| 18S.1.5.F.PCM.ME | Mawson Escarpment | 68,42333333 | -73,30916667 |
| 18S.1.5.H.PCM.ME | Mawson Escarpment | 68,36805556 | -73,32805556 |
| 18S.1.6.D.PCM.ME | Mawson Escarpment | 68,14138889 | -72,92361111 |
| 18S.1.6.E.PCM.ME | Mawson Escarpment | 68,46555556 | -73,30027778 |
| 18S.1.6.F.PCM.ME | Mawson Escarpment | 68,40555556 | -73,30805556 |
| 18S.1.6.G.PCM.ME | Mawson Escarpment | 68,48722222 | -73,30555556 |
| 18S.1.6.H.PCM.ME | Mawson Escarpment | 68,37555556 | -73,33 |
| 18S.1.7.D.PCM.ME | Mawson Escarpment | 68,30138889 | -73,32611111 |
| 18S.1.7.E.PCM.ME | Mawson Escarpment | 68,35361111 | -73,31583333 |
| 18S.1.7.F.PCM.ME | Mawson Escarpment | 68,42666667 | -73,31638889 |
| 18S.1.7.G.PCM.ME | Mawson Escarpment | 68,36277778 | -73,32333333 |
| 18S.1.7.H.PCM.ME | Mawson Escarpment | 68,38444444 | -73,32694444 |
| 18S.1.8.D.PCM.ME | Mawson Escarpment | 68,34111111 | -73,3225 |
| 18S.1.8.E.PCM.ME | Mawson Escarpment | 68,38472222 | -73,31055556 |
| 18S.1.8.F.PCM.ME | Mawson Escarpment | 68,42166667 | -73,31083333 |
| 18S.1.8.G.PCM.ME | Mawson Escarpment | 68,36055556 | -73,32333333 |
| 18S.1.8.H.PCM.ME | Mawson Escarpment | 68,38055556 | -73,31777778 |
| 18S.1.9.D.PCM.ME | Mawson Escarpment | 68,42333333 | -73,30555556 |
| 18S.1.9.E.PCM.ME | Mawson Escarpment | 68,39638889 | -73,2775 |
| 18S.1.9.F.PCM.ME | Mawson Escarpment | 68,38944444 | -73,31444444 |
| 18S.1.9.G.PCM.ME | Mawson Escarpment | 68,5 | -73,30972222 |
| 18S.1.9.H.PCM.ME | Mawson Escarpment | 68,37222222 | -73,31388889 |
| 18S.2.1.A.PCM.ME | Mawson Escarpment | 68,35555556 | -73,31583333 |
| 18S.2.1.B.PCM.ME | Mawson Escarpment | 66,46972222 | -73,01972222 |
| 18S.2.10.A.PCM.ME | Mawson Escarpment | 66,445 | -73,01194444 |
| 18S.2.2.A.PCM.ME | Mawson Escarpment | 68,35833333 | -73,31416667 |
| 18S.2.2.B.PCM.ME | Mawson Escarpment | 68,41638889 | -73,48416667 |
| 18S.2.2.H.PCM.ME | Mawson Escarpment | 68,33944444 | -73,32111111 |
| 18S.2.3.A.PCM.ME | Mawson Escarpment | 68,49055556 | -73,30888889 |
| 18S.2.4.A.PCM.ME | Mawson Escarpment | 68,24444444 | -73,29861111 |
| 18S.2.4.B.PCM.ME | Mawson Escarpment | 66,79472222 | -74,29027778 |
| 18S.2.5.A.PCM.ME | Mawson Escarpment | 68,24777778 | -73,30888889 |
| 18S.2.5.B.PCM.ME | Mawson Escarpment | 65,60944444 | -73,39638889 |
| 18S.2.6.A.PCM.ME | Mawson Escarpment | 68,31777778 | -73,31972222 |
| 18S.2.6.B.PCM.ME | Mawson Escarpment | 68,15 | -73,19611111 |
| 18S.2.7.A.PCM.ME | Mawson Escarpment | 68,04166667 | -72,81944444 |
| 18S.2.8.A.PCM.ME | Mawson Escarpment | 68,47 | -73,63777778 |
| 18S.2.9.A.PCM.ME | Mawson Escarpment | 68,52944444 | -73,64388889 |
| 18S.2.1.C.PCM.LT | Lake Terrasovoje | 67,88055556 | -70,54194444 |
| 18S.2.1.D.PCM.LT | Lake Terrasovoje | 67,89666667 | -70,53083333 |
| 18S.2.1.E.PCM.LT | Lake Terrasovoje | 67,94694444 | -70,55916667 |
| 18S.2.1.F.PCM.LT | Lake Terrasovoje | 67,79777778 | -70,525 |
| 18S.2.1.G.PCM.LT | Lake Terrasovoje | 68,2125 | -70,82833333 |
| 18S.2.1.H.PCM.LT | Lake Terrasovoje | 68,28472222 | -73,41277778 |
| 18S.2.10.B.PCM.LT | Lake Terrasovoje | 68,01638889 | -70,51222222 |
| 18S.2.10.C.PCM.LT | Lake Terrasovoje | 68,01388889 | -70,53305556 |
| 18S.2.10.D.PCM.LT | Lake Terrasovoje | 67,88972222 | -70,52361111 |
| 18S.2.10.E.PCM.LT | Lake Terrasovoje | 67,98361111 | -70,54888889 |
| 18S.2.10.F.PCM.LT | Lake Terrasovoje | 68,07111111 | -70,84472222 |
| 18S.2.10.G.PCM.LT | Lake Terrasovoje | 68,80805556 | -70,42416667 |
| 18S.2.11.B.PCM.LT | Lake Terrasovoje | 67,82555556 | -70,52722222 |
| 18S.2.11.C.PCM.LT | Lake Terrasovoje | 69,00805556 | -70,53388889 |
| 18S.2.11.D.PCM.LT | Lake Terrasovoje | 67,99305556 | -70,54305556 |
| 18S.2.11.E.PCM.LT | Lake Terrasovoje | 67,96916667 | -70,54333333 |
| 18S.2.11.F.PCM.LT | Lake Terrasovoje | 68,0425 | -70,84472222 |
| 18S.2.2.C.PCM.LT | Lake Terrasovoje | 67,83305556 | -70,53027778 |
| 18S.2.2.D.PCM.LT | Lake Terrasovoje | 68,00222222 | -70,52888889 |
| 18S.2.2.E.PCM.LT | Lake Terrasovoje | 67,93944444 | -70,55694444 |
| 18S.2.2.F.PCM.LT | Lake Terrasovoje | 67,81083333 | -70,52666667 |
| 18S.2.2.G.PCM.LT | Lake Terrasovoje | 68,02 | -70,80861111 |
| 18S.2.3.C.PCM.LT | Lake Terrasovoje | 68,05166667 | -70,51722222 |
| 18S.2.3.D.PCM.LT | Lake Terrasovoje | 67,92722222 | -70,51694444 |
| 18S.2.3.E.PCM.LT | Lake Terrasovoje | 67,91611111 | -70,54805556 |
| 18S.2.3.F.PCM.LT | Lake Terrasovoje | 67,82305556 | -70,52722222 |
| 18S.2.3.G.PCM.LT | Lake Terrasovoje | 68,00861111 | -70,80916667 |
| 18S.2.4.C.PCM.LT | Lake Terrasovoje | 68,04222222 | -70,52138889 |
| 18S.2.4.D.PCM.LT | Lake Terrasovoje | 67,89027778 | -70,53138889 |
| 18S.2.4.E.PCM.LT | Lake Terrasovoje | 67,77416667 | -70,53972222 |
| 18S.2.4.F.PCM.LT | Lake Terrasovoje | 67,86777778 | -70,52277778 |
| 18S.2.4.G.PCM.LT | Lake Terrasovoje | 68,17666667 | -70,82916667 |
| 18S.2.5.C.PCM.LT | Lake Terrasovoje | 67,87222222 | -70,52666667 |
| 18S.2.5.D.PCM.LT | Lake Terrasovoje | 68,00361111 | -70,52638889 |
| 18S.2.5.F.PCM.LT | Lake Terrasovoje | 68,03416667 | -70,54027778 |
| 18S.2.5.G.PCM.LT | Lake Terrasovoje | 64,60361111 | -70,42944444 |
| 18S.2.6.C.PCM.LT | Lake Terrasovoje | 67,87361111 | -70,52388889 |
| 18S.2.6.D.PCM.LT | Lake Terrasovoje | 67,99777778 | -70,52027778 |
| 18S.2.6.E.PCM.LT | Lake Terrasovoje | 67,85805556 | -70,54583333 |
| 18S.2.6.F.PCM.LT | Lake Terrasovoje | 68,04027778 | -70,55027778 |
| 18S.2.7.C.PCM.LT | Lake Terrasovoje | 68,00388889 | -70,5175 |
| 18S.2.7.D.PCM.LT | Lake Terrasovoje | 67,91583333 | -70,51833333 |
| 18S.2.7.E.PCM.LT | Lake Terrasovoje | 67,98805556 | -70,58388889 |
| 18S.2.7.F.PCM.LT | Lake Terrasovoje | 68,17722222 | -70,80916667 |
| 18S.2.8.B.PCM.LT | Lake Terrasovoje | 68,02944444 | -70,51277778 |
| 18S.2.8.C.PCM.LT | Lake Terrasovoje | 67,87861111 | -70,52194444 |
| 18S.2.8.D.PCM.LT | Lake Terrasovoje | 67,91861111 | -70,53333333 |
| 18S.2.8.E.PCM.LT | Lake Terrasovoje | 67,90722222 | -70,54611111 |
| 18S.2.8.F.PCM.LT | Lake Terrasovoje | 68,06777778 | -70,81194444 |
| 18S.2.8.G.PCM.LT | Lake Terrasovoje | 66,895 | -70,56916667 |
| 18S.2.9.C.PCM.LT | Lake Terrasovoje | 67,90611111 | -70,54361111 |
| 18S.2.9.D.PCM.LT | Lake Terrasovoje | 67,98694444 | -70,54055556 |
| 18S.2.9.E.PCM.LT | Lake Terrasovoje | 67,98138889 | -70,57527778 |
| 18S.2.9.F.PCM.LT | Lake Terrasovoje | 68,06472222 | -70,83472222 |
| 18S.2.9.G.PCM.LT | Lake Terrasovoje | 68,3825 | -70,80944444 |
