## Supplementary material for "Antarctic biodiversity predictions through substrate qualities and environmental DNA": Web Table 3

**Web Table 3:** Project sequencing effort and obtained read to yield unfiltered data (including controls), across all sequencing runs.

| Run | Input | Filtered | Denoised | Non-chimeric | Libraries | Avg. Seq./Lib. |
| --- | --- | --- | --- | --- | --- | --- |
| A | 10,055,133 | 9,771,027 | 9,759,212 | 9,667,648 | 92 | 105,083 |
| B | 10,061,999 | 9,872,492 | 9,856,147 | 9,725,926 | 91 | 106,818 |
|  | 20,117,132 | 19,643,519 | 19,615,359 | 19,393,574 | 183 | 105,975 |
