## Supplementary material for "Antarctic biodiversity predictions through substrate qualities and environmental DNA": Web Table 7

**Web Table 2:** Signs of coefficient estimates for each predictor resulting from lasso logistic regression. Signs indicate changes at Mawson Escarpment and Mount Menzies in comparison to the reference location (Lake Terrasovoje). X-ray spectral peaks of certain wavelength are compounds of mineral group as indicated by asterisk (*).

| Phylum | Potassium | Sulphur | Conductivity | pH of CaCl2 | ATP fluorescence | Feldspar | Titanite | Pyroxene / Amphibole / Garnet * | Micas | Dolomite | Kaolin / Chlorite* | Calcite | Chlorite | Slope |
| --- | --- | --- | --- | --- | --- | --- | --- | --- | --- | --- | --- | --- | --- | --- |
| Arthropoda |  |  |  |  |  | + |  |  | - | - |  |  |  |  |
| Ascomycota |  |  |  | - | + |  | + |  |  |  |  |  |  |  |
| Bacillariophyta |  |  | - |  |  |  |  | - | - |  | + |  |  | - |
| Basidiomycota |  | - |  | - |  | + | + |  |  | + |  |  | - |  |
| Cercozoa |  | - |  | - |  |  |  |  |  |  |  | - | - |  |
| Chlorophyta | - |  |  | - |  |  |  |  |  |  | + | + |  | - |
| Chytridiomycota |  |  | - | - | - |  |  | - | - | + | + |  |  |  |
| Ciliophora |  | - | - |  |  |  |  | - | - |  | + |  |  |  |
| Nematoda |  | - | - | + |  |  |  |  |  |  |  |  |  |  |
| Rotifera | - |  |  | - |  | + | + |  |  |  |  |  |  |  |
| Streptophyta |  |  |  |  |  |  |  |  |  |  | - |  |  |  |
