## Supplementary material for "Antarctic biodiversity predictions through substrate qualities and environmental DNA": Web Text 1

Antarctic biodiversity predictions through substrate qualities and environmental DNA – Online Supplemental Material

### On sequence data generation

- Czechowski *et al.* (2016a, b) describe sequence data generation in high detail.

### On data filtering and taxonomic assignments

- Our eDNA data reprocessing started with 16,524,031 sequences, which collectively represented 2,656 eukaryotes, as well as 5,441 non-eukaryotes (i.e., bacterial, archean or undetermined) ASVs. Unfiltered sample mean (min. / med. / max.) coverage was 107,299 reads (1 / 108, 773 / 692, 828), unfiltered ASV mean (min. / med. / max.) coverage was 2,041 reads (2 / 70 / 1, 491, 686). After application of package *decontam* (Davis *et al.* 2018) some ASVs remained in control reactions, prior to their subtraction from field data: one ASV of indefinable origin in the extraction blanks, and 51 bacterial and eukaryote ASVs in the amplification blanks.
- Please refer to Web Tables 4 and 5 for alignment qualities.
- Example references for the five highest-covered species among our data are:
  - *Acanthothecis fontana*: Muscavitch and Lendemer (2016)
  - *Coccomyxa* sp.: Blanc *et al.* (2012)
  - *Mrakia frigida:* Xin and Zhou (2007)
  - *Pseudochilodonopsis quadrivacuolata*: Qu *et al.* (2015)
  - *Scottnema lindsayae*:	Velasco-Castrillón *et al.* (2014)
  - *Embata laticeps*: (Myers (1931)
  - *Mesobiotus furciger*: Velasco-Castrillón *et al.* (2014)

### On geographic distribution of taxonomic assignments

- We investigated the geographic distribution of the significant 173 species assignments by using R package *spocc* (Chamberlain 2021). We cleaned the species names retrieved from NCBI using regular-expression matching, and submitted them as queries to BISON (United States Federal Resource for Biological Occurrence Data ­– <https://bison.usgs.gov>), GBIF (Global Biodiversity Information Facility – <https://www.gbif.org> ), and iNaturalist (<https://www.inaturalist.org>), requesting up to 10 georeferenced occurrence records for each query. We obtained a total of 778 records (53 from BISON, 523 from GBIF, and 202 from iNatuarlist). Successfully georeferenced species are listed by continent in Web Table 6.
- For 173 queries, occurrence records could be obtained from public repositories for the 66 taxonomic assignments of:
  - *Amanita brunnescens, Agrocybe rivulosa, Hebeloma crustuliniforme, Rhodocybella rhododendri, Dictyonema sericeum, Mycetinis alliaceus, Gloiocephala aquatica, Auricularia auricula-judae, Suillus lakei, Leucogyrophana romellii, Rhizoctonia solani, Limonomyces roseipellis, Rogersella griseliniae, Hymenochaete cruenta, Resinicium friabile, Resinicium mutabile, Rigidoporus corticola, Donkia pulcherrima, Phanerochaete chrysosporium, Stereum rugosum, Buckleyzyma aurantiaca, Cerinomyces canadensis, Glaciozyma antarctica, Rhodotorula glutinis, Rhodotorula mucilaginosa, Cystofilobasidium infirmominiatum Mrakia frigida, Udeniomyces puniceus, Vishniacozyma dimennae, Syzygospora alba, Apiotrichum domesticum, Tritirachium egenum, Ustilago bromivora, Oophila amblystomatis, Haematococcus lacustris, Rusalka fusiformis, Pascherina tetras, Desmococcus olivaceus, Catena viridis, Closteriopsis acicularis, Koliella spiculiformis, Prasiola crispa, Chloroidium saccharophilum, Phyllosiphon arisari, Pseudocyrtolophosis alpestris, Mykophagophrys terricola, Pelagodileptus trachelioides, Apobryophyllum schmidingeri, Furgasonia blochmanni, Nassula labiata, Stokesia vernalis, Cohnilembus verminus, Pseudocohnilembus persalinus, Cristigera media, Opercularia microdiscum, Chilodonella acuta, Pseudochilodonopsis mutabilis, Tachysoma pellionellum, Eucephalobus oxyuroides, Scottnema lindsayae, Pratylenchus thornei, Pelodera teres, Milnesium tardigradum, Diphascon pingue, Macrobiotus hufelandi, Echiniscus jenningsi*
- For 173 queries, occurrence records could not be obtained via public repositories for the 107 taxonomic assignments of:
  - *Porpolomopsis* sp.*, Marasmius* sp.*, Laccaria* sp.*, Proceropycnis pinicola, Bannoa* sp.*, Meira* sp.*, Golubevia* sp.*, Yamadamyces* sp.*, Leucosporidium* sp.*, Chrysozyma* sp.*, Aecidium kalanchoes, Goffeauzyma iberica, Goffeauzyma metallitolerans, Holtermannia* sp.*, Holtermanniella nyarrowii, Fonsecazyma* sp.*, Genolevuria* sp.*, Derxomyces* sp.*, Hannaella pagnoccae, Hannaella* sp.*, Vishniacozyma carnescens, Vishniacozyma peneaus, Vishniacozyma* sp.*, Cryptococcus* sp.*, Phaeotremella* sp.*, Apiotrichum loubieri, Apiotrichum wieringae, Trichosporon* sp.*, Wallemia* sp.*, Diplosphaera chodatii, Pleurococcus* sp.*, Chloromonas fonticola, Chloromonas nivalis, Chloromonas* sp.*, Chloromonas* sp.*, Chloromonas subdivisa, Rhysamphichloris curta, Characium perforatum, Hafniomonas reticulata, Sanguina nivaloides, Dictyococcus* sp.*, Mychonastes* sp.*, Follicularia texensis, Scenedesmus* sp.*, Tetradesmus incrassatulus, Sphaerocystis* sp.*, Monomastix minuta, Monomastix* sp.*, Monomastix* sp.*, Apatococcus* sp.*, Dictyosphaerium* sp.*, Micractinium* sp.*, Micractinium* sp.*, Micractinium* sp.*, Micractinium* sp.*, Coenochloris* sp.*, Stichococcus antarcticus, Stichococcus* sp.*, Stichococcus* sp.*, Prasiola* sp.*, Chloroidium angustoellipsoideum, Coccomyxa* sp.*, Pseudostichococcus monallantoides, Rhexinema paucicellulare, Sarcinofilum mucosum, Ottowphyra dragescoi, Monodinium* sp.*, Dileptus jonesi, Enchelys megaspinata, Lacrymaria* sp.*, Phialina vertens, Acaryophrya* sp.*, Amphileptus* sp.*, Apocolpodidium etoschense, Nassula* sp.*, Naxella paralucida, Tetrahymena* sp.*, Trichodinella* sp.*, Platynematum salinarum, Rhabdostyla* sp.*, Vaginicola* sp.*, Biggaria bermudensis, Pseudochilodonopsis quadrivacuolata, Pseudochilodonopsis* sp.*, Microdysteria decora, Levicoleps taehwae, Halodinium verrucatum, Hemiurosomoida longa, Oxytricha paragranulifera, Urosomoida agilis, Lamtostyla ovalis, Amphisiella pulchra, Onychodromus quadricornutus, Keronopsis helluo, Urospinula succisa, Parabistichella variabilis, Paraholosticha pannonica, Pseudourostyla nova, Halomonhystera* sp.*, Halicephalobus* sp.*, Allodorylaimus* sp.*, Discolaimus, Acutuncus antarcticus, Mesobiotus furciger, Minibiotus gumersindoi, Minibiotus* sp.*, Pseudechiniscus titianae*
